# Pathway and mechanism of tubulin folding mediated by TRiC/CCT conjugated with its ATPase cycle revealed by cryo-EM

**DOI:** 10.1101/2022.08.08.503128

**Authors:** Caixuan Liu, Mingliang Jin, Shutian Wang, Wenyu Han, Qiaoyu Zhao, Yifan Wang, Cong Xu, Lei Diao, Yue Yin, Chao Peng, Lan Bao, Yanxing Wang, Yao Cong

## Abstract

The eukaryotic chaperonin TRiC/CCT assists the folding of about 10% of cytosolic proteins through an ATP-driven conformational cycle, and the essential cytoskeleton protein tubulin is the obligate substrate of TRiC. Here, we present an ensemble of cryo-EM structures of human TRiC throughout its ATPase cycle, with three of them revealing endogenously engaged tubulin in different folding stages. Our structural and XL-MS analyses suggested a gradual upward translocation and stabilization of tubulin within the TRiC chamber accompanying TRiC ring closure. Remarkably, in the closed TRiC-tubulin-S3 map resolved to 3.1-Å-resolution, we captured a near-natively folded tubulin. We found the near-natively folded tubulin engaging through its N and C domains mainly with the A and I domains of the CCT3/6/8 subunits through electrostatic and hydrophilic interactions, while the tubulin I domain was found to remain dynamic. Moreover, we also showed the potential role of TRiC C-terminal tails in substrate stabilization and folding. Our study delineates the pathway and molecular mechanism of TRiC-mediated tubulin folding conjugated with TRiC ATPase cycle, and may also inform the design of therapeutic agents targeting TRiC-tubulin interactions.

## Introduction

Chaperonins are protein folding nanomachines that play an essential role in maintaining cellular homeostasis and are important in all kingdoms of life, and their dysfunction is closely related to cancer and neurodegenerative diseases^1–3^. Chaperonins provide a cage-like environment for proteins to fold in isolation, unimpaired by aggregation, and in some cases actively modulate the folding pathway of the encapsulated protein^4–7^. The eukaryotic group II chaperonin TRiC/CCT assists the folding of ∼10% of cytosolic proteins, including the key cytoskeletal proteins actin and tubulin, cell cycle regulator CDC20, and VHL tumor suppressor^8–15^. TRiC is the most complex chaperon system identified to date. It has a double-ring structure, and each ring consists of eight paralogous subunits (namely CCT1-CCT8) arranged in a specific order^16–21^. Each TRiC subunit consists of three domains: the substrate-recognition apical domain (A domain) and the ATP-binding equatorial domain (E domain) linked by the intermediate domain (I domain). TRiC-mediated substrate folding is closely related to its ATP-driven conformational cycle^22–25^. TRiC has been proved to display subunit specificity in the complex assembly, ATP consumption and ring closure^16, 19, 26, 27^.

In the past two decades, constant efforts have been made to capture the TRiC-substrate binary complexes so as to reveal the mechanism of TRiC-assisted substrate folding for diverse substrates^7, 9, 11, 15, 28–32^. Still, there remains lacking of the atomic-resolution structural details revealing the dynamic process of TRiC-mediated substrate folding throughout its ATP-driven conformational cycle, due to the relative low binding efficiency between TRiC and the pretreated unfolded substrates (usually involving applying urea or GuHCl *in vitro*) and due to the potential conformational and compositional heterogeneity of the binary complex^33^.

Tubulin, the building block of microtubule, is the *in vivo* obligate substrate of TRiC^15, 34–37^. Highly conserved α- and β-tubulin heterodimers assemble into dynamic microtubules^38^, which are ubiquitous cytoskeletal polymers essential for the life of all eukaryotic cells. Microtubules are involved in a wide range of cellular functions, from cell motility and intracellular transport to, most fundamentally, cell division^36, 39–43^. Tubulin (∼50 kDa) contains three domains: the GTP-binding N domain, the I domain containing the site that binds the chemotherapeutic Taxol, and the C domain responsible for interacting with microtubule-binding proteins. Previous studies reported structures of the TRiC-tubulin complex in the apo or ATP-binding state at relatively low resolution (∼25 Å) or intermediate resolution (5.5 Å) determined using cryo-electron microscopy (cryo-EM) or X-ray crystallography^9, 15, 32^. However, a complete picture with molecular details of the TRiC-directed tubulin folding driven by ATP binding and hydrolysis remains obscure. Moreover, it has been reported that the small molecule I-Trp can disrupt constitutively associated β-tubulin/TRiC, causing severe cell apoptosis, indicating that targeting the TRiC/tubulin complex could serve as a novel chemotherapeutic strategy^44, 45^.

To capture the TRiC-directed tubulin folding pathway, we determined an ensemble of cryo-EM structures of human TRiC (hTRiC) throughout its ATP-driven conformational cycle. Three of these structures showed endogenously bound tubulin. Strikingly, in the closed-state TRiC-tubulin-S3 structure, we captured a near-natively folded tubulin and disclosed atomic details of the interaction between tubulin and TRiC. We also found that the C- and N-terminal tails (abbreviated as C-/N-termini) of TRiC play distinct roles in substrate stabilization/folding and TRiC allosteric coordination, respectively, with the N-terminus of CCT8 possibly contributing to tubulin folding in the opposite ring. Collectively, our study captured a thorough picture of the pathway and molecular mechanism of TRiC-mediated tubulin folding conjugated with the TRiC ATPase cycle.

## Results

### Cryo-EM structure of TRiC in complex with tubulin

We purified endogenous human TRiC from HEK293F cells (Extended Data Fig. 1a; hereafter, “TRiC” refers to human TRiC unless otherwise noted), which was validated by sodium dodecyl sulfate-polyacrylamide gel electrophoresis (SDS-PAGE) and mass spectrometry (MS) analyses (Fig. 1a and Supplementary Table 1). The result of NADH-coupled enzymatic assay further validated the ATPase activity of the purified TRiC (Extended Data Fig. 1b). Moreover, our SDS-PAGE analysis also indicated the presence of an extra associated protein at ∼50 kDa (Fig. 1a), which was suggested to be mainly β-tubulin (TUBB5) by the MS analysis (Supplementary Table 1). The PSM value from the MS analysis, indicating the relative abundance of a certain protein^46^, showed the abundance of β-tubulin was comparable to that of individual TRiC subunit. The existence of tubulin was further confirmed by native electrophoresis combined with western blot and chemical cross-linking-coupled mass spectrometry (XL-MS) analysis (Fig. 1b-c, Supplementary Table 2). Our XL-MS analysis detected five cross-links between tubulin and CCT3/4/6/8 subunits (Fig. 1c). Taken together, these data indicated a co-existence of TRiC with tubulin.

**Fig. 1.**
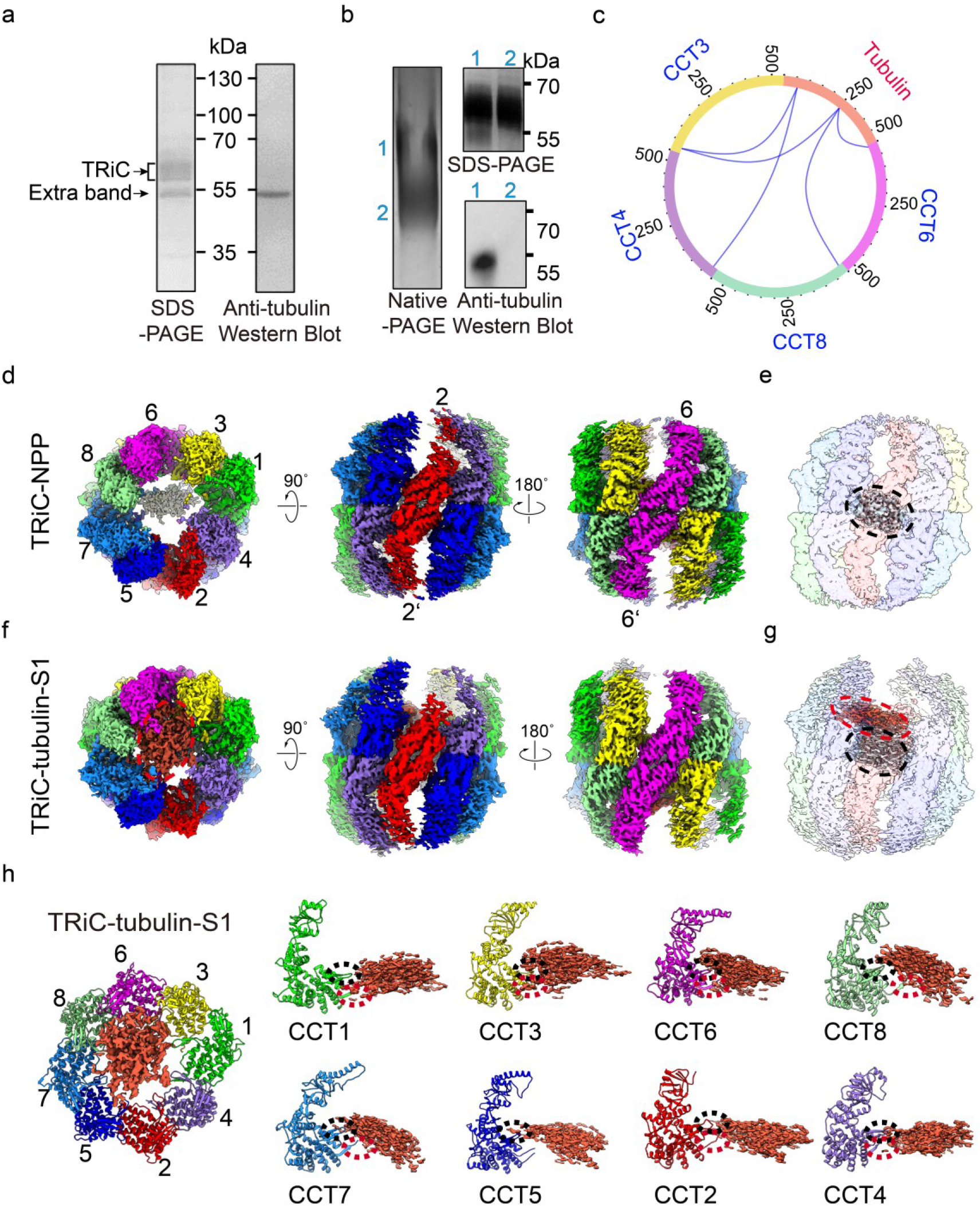
Cryo-EM structures of the endogenously purified TRiC with associated tubulin. **a**, Sodium dodecyl sulfate-polyacrylamide gel electrophoresis (SDS-PAGE) of the endogenous human TRiC purified from HEK293F cells. This gel suggested the presence of an extra associated protein at ∼50 kDa, which was proved to be tubulin by Western blot. **b**, Native gel analysis of TRiC, showing two bands, labeled as 1 and 2, which were excised and loaded into SDS-PAGE. SDS-PAGE analysis suggested that both bands contain TRiC oligomer, and there is an additional band in lane 1, which contained tubulin, as shown by Western blot. **c**, XL-MS analysis of the endogenously purified TRiC with associated tubulin. Identified cross-linked TRiC-tubulin contacts are shown as blue lines. **d,** End-on and side views of TRiC-NPP, with the different subunits in different colors and labeled. This subunit color scheme is followed in subsequent figures. **e,** The unstructured N- and C-terminal tail density (in grey, indicated by a black dashed ellipsoid) of TRiC subunits located between the two equators in TRiC-NPP (transparent density). **f-g,** Cryo-EM map of TRiC-tubulin-S1 (**f**), revealing extra density (shown in red and indicated by a red dashed ellipsoid) within the cis-ring chamber of TRiC (**g**). **h**, Overall binding location of tubulin (red density) in the cis-ring of TRiC-tubulin-S1 (colored ribbon) (left), and the association of tubulin with the E domain of every TRiC subunit, including the C-terminus (indicated by dotted red ellipsoid) and stem loop (dotted black ellipsoid) (right panels).

We then performed cryo-EM study on the endogenously purified TRiC sample (Extended Data Fig. 1c). Two cryo-EM maps, including an open state TRiC with an empty chamber, and another TRiC map also in the open conformation but displaying an extra density in one chamber, were determined at 3.1- and 4.1-Å-resolution, respectively (Fig. 1d-g, and Extended Data Fig. 2). Interestingly, although we performed buffer exchange multiple times to remove remaining nucleotide in the final stage of purification, CCT3/6/8 subunits appeared to have bound nucleotide density in both maps (Extended Data Fig. 3e-f), which were suggested to be ADP according to an ADP/ATP ratio assay (Extended Data Fig. 3g), in line with our previous report on the yeast TRiC system^19^. Accordingly, these two open TRiC maps are also in the “nucleotide partially preloaded” (NPP) state. We then denoted the free open TRiC map as TRiC-NPP. The NPP state is equivalent to the “apo” state in typical conformational cycle studies of ATPase chaperonins^24, 47, 48^, mimicking the starting point in the conformational cycle of the chaperonin itself.

Our TRiC-NPP map showed a conformation overall similar to that of the available substrate-free human TRiC cryo-EM map also in the open state at 7.7-Å-resolution, with determined subunit assignment^49^. We then fit the TRiC-NPP map to their map (Extended Data Fig. 4a), both displaying the following characteristic features known for open-state TRiC: (1) CCT1 being the most outward tilted subunit, a feature common for yeast, bovine, and human TRiCs^7, 16, 19, 49^; (2) the A domain of CCT2 being quite disordered (Fig. 1d), also observed in open bovine TRiC^7, 24^; (3) each ring displaying a tetramer-of-dimer pattern as in the open bovine TRiC^24^; (4) the largest gap existing between CCT1 and CCT4^50^ (Extended Data Fig. 4b). These features allowed us to assign the subunits for the TRiC-NPP map. We then built an atomic model for TRiC-NPP (Extended Data Fig. 3a-d). Further inspection of the TRiC-NPP structure revealed the characteristic V476-K484 insertion in the E domain of CCT1 (Extended Data Fig. 4c-d), corroborating our subunit assignment for this map.

Moreover, our TRiC-NPP map showed a chunk of density between the two equators blocking the two chambers (Fig. 1e), with this density symmetrically contacting the N-/C-terminal extensions of CCT5/7 and CCT5’/7’ from both rings of TRiC (Extended Data Fig. 4e). Hence, we assigned the density as the unstructured N- and C-terminal tails of TRiC subunits, also observed in the available human and bovine TRiC open-state maps^7, 49^ (Extended Data Fig. 4f-g). Moreover, the A/I domains of CCT2 hemisphere subunits (including CCT4/2/5/7)^21, 28^ appeared less well resolved as in the bovine and human open-state TRiC structures^7, 24, 28, 49^, indicating an intrinsic dynamic nature in these regions. Indeed, our 3D variability analysis (3DVA) using cryoSPARC^51^ suggested the A/I domains of CCT1/4/2/5/7 to be overall relatively dynamic—and strikingly, with those of CCT7/5/1 showing continuous outward/inward tilting motions of up to ∼23°/6°/5°, respectively (Extended Data Fig. 4h and Supplementary Video 1).

Importantly, for the other map derived from the same dataset (38.4% of the population, Fig. 1f-g and Extended Data Fig. 2a), we attributed the captured extra density in the cis-ring chamber to the trapped β-tubulin co-purified with TRiC, based on our biochemical and MS data (Fig. 1a-c, Extended Data Fig. 4i and Supplementary Table 1-2). We denoted this map as TRiC-tubulin-S1, which was also in the NPP state with the consecutive CCT3/6/8 subunits loaded with nucleotides in both rings (Extended Data Fig. 3f). This demonstrated that TRiC can engage with substrate in the nucleotide partially loaded NPP state. A previous crystal structure of bovine TRiC with bound tubulin also revealed two nucleotides per-ring in non-consecutive subunits^15^. The TRiC conformation in the S1 map was observed to resemble that of our TRiC-NPP structure (Extended Data Fig. 4j), we then followed the same subunit assignment. Overall, the substrate density appeared to associate with the E domain of every TRiC subunit, mainly with the C-terminus and stem loop of these subunits (Fig. 1h). The bovine TRiC-tubulin crystal structure also revealed that the tubulin forms contact with the stem loops of three TRiC subunits^15^.

### Tubulin translocation within TRiC chamber accompanying TRiC conformational cycle

To further capture the TRiC-directed tubulin folding process conjugated with TRiC ATPase cycle, we performed cryo-EM study on TRiC in the presence of 1 mM ATP-AlFx (Extended Data Fig. 1d), an ATP-hydrolysis-transition state analog that can trigger TRiC ring closure^22, 23, 25^. In the resulting dataset, the majority (65.4%) of the particles were in the closed conformation and resolved to a resolution of 2.9 Å, while the remaining (34.6%) particles were in the open conformation and resolved to a resolution of 4.2 Å (Fig. 2a and Extended Data Fig. 5a-c). In the open state, obvious substrate density was observed in the cis-ring, and we denoted this map as TRiC-tubulin-S2. For the closed state, through focused classification in the chamber region, we obtained two maps: one map at a resolution of 3.1 Å showed tubulin density in the cis-ring chamber (termed TRiC-tubulin-S3), and the other map at a resolution of 3.2 Å showed no substrate in the chamber, but a central tail density in an orientation different from that in the open state (denoted as TRiC-ADP-AlFx, discussed below) (Fig. 2c-g, Extended Data Fig. 5a-b, d-f). Moreover, we performed XL-MS analysis on the ATP-AlFx presented TRiC sample, and detected 17 cross-links between tubulin and TRiC subunits (Fig. 2h and Supplementary Table 3), further substantiating the notion that the captured substrate was indeed tubulin.

**Fig. 2.**
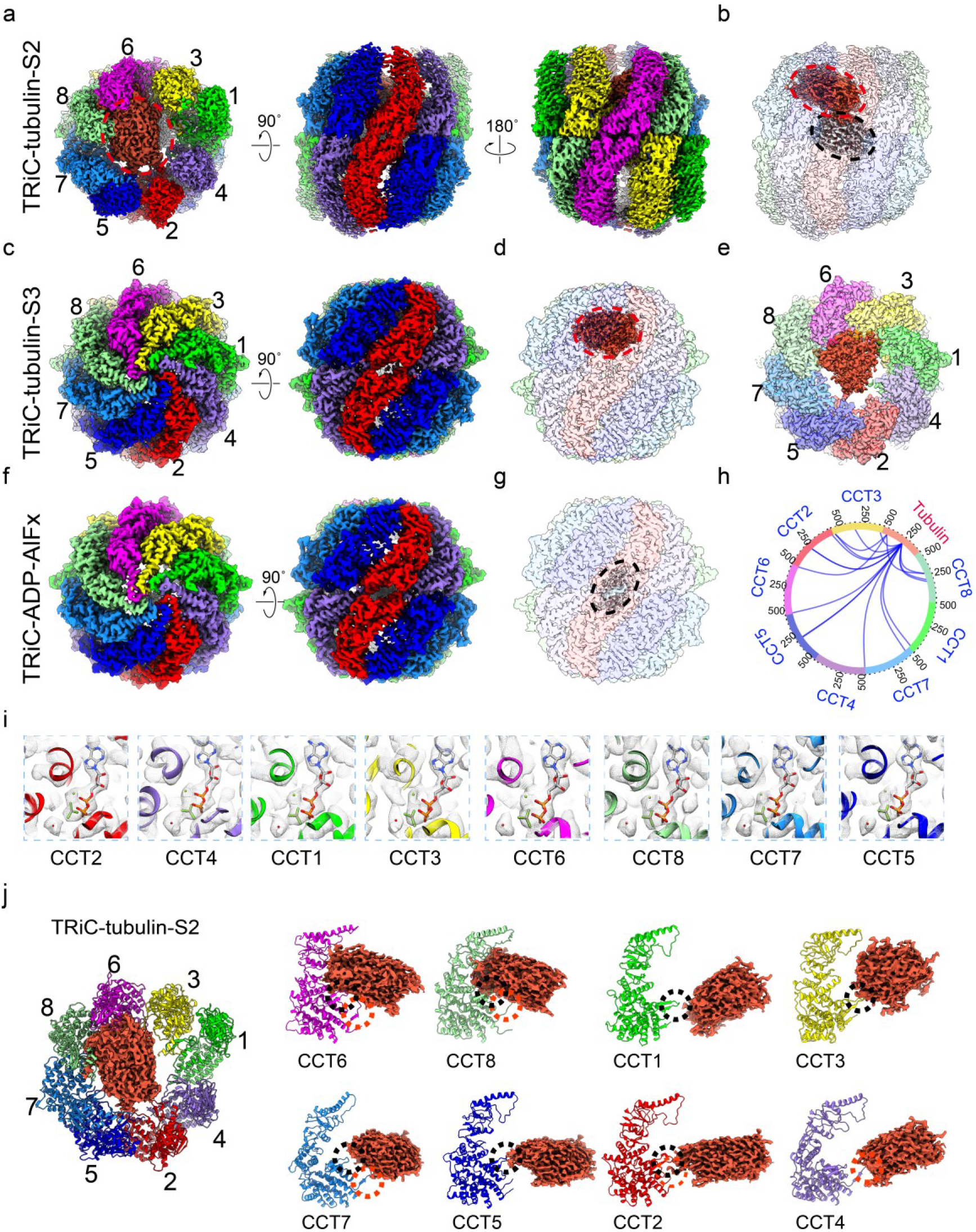
Cryo-EM structures of TRiC-tubulin in the presence of ATP-AlFx. **a-b,** Cryo-EM map of TRiC-tubulin-S2 in the open ATP-binding state (**a**), revealing a tail density (in grey) between the two equators and a tubulin density (in red) in the cis-ring of TRiC (transparent density) (**b**). The tail density contacts the tubulin density. **c-d,** Cryo-EM map of TRiC-tubulin-S3 in the closed ATP-hydrolysis transition state (**c**), revealing tubulin density in the cis-ring chamber of TRiC (**d**). **e**, Top view of the TRiC-tubulin-S3 map with TRiC A domains omitted for easier visualization, showing the bound tubulin density, and indicating tubulin mainly associate with CCT6 hemisphere subunits (CCT1/3/6/8). **f-g,** Cryo-EM map of TRiC-ADP-AlFx (**f**), revealing a C-termini mass (black dashed ellipsoid) (**g**), related to Extended Data Fig. 5f. **h**, XL-MS analysis of TRiC with endogenously associated tubulin in the presence of ATP-AlFx. **i**, Portions of the TRiC-tubulin-S3 map in the nucleotide pocket region. All of the subunits bound with ADP-AlFx (stick model) and a magnesium ion (green ball), as well as a water molecule (red ball) in an attacking position, suggesting that all eight subunits are in the ATP-hydrolysis transition state. **j**, Overall binding of tubulin (red density) in the cis-ring of the open TRiC-tubulin-S2. Tubulin was observed to contact all three domains of CCT6/8 and loosely contact the E domains, including the C terminus (dotted red ellipsoid) and stem loop (dotted black ellipsoid), of the remaining subunits.

The overall TRiC conformation of the open-state S2 map was observed to be similar to that of the TRiC-NPP map (Extended Data Fig. 5g). We then followed the subunit assignments of TRiC-NPP to assign those of S2. For the closed-state S3 map, a characteristic kink feature in the CCT6 α-helical protrusion H8 and the unique insertions in CCT1/4/6 were clearly visualized (Extended Data Fig. 6), facilitating the subunit assignment in the maps of the closed TRiC with similar features. We then built an atomic model for each of the S2, S3 and TRiC-ADP-AlFx maps (Extended Data Fig. 7a-f). We found that for these three structures, all subunits from both rings bound nucleotides (Extended Data Fig. 7g-i). For the S2 map, the nucleotide density was fitted well by the ATP structure (Extended Data Fig. 7j), indicating S2 was in the ATP-binding state. For the closed S3 and TRiC-ADP-AlFx structures, with their E-domain local resolution reached ∼2.8 to 2.9 Å (Extended Data Fig.5d-e), the nucleotide densities for all subunits matched ADP-AlFx and a magnesium ion very well, in addition to a water molecule attacking the γ-phosphate of nucleotide, involved in the formation of the ATP hydrolysis reaction center (Fig. 2i and Extended Data Fig. 7k). These features suggested the two closed-state maps were in the ATP hydrolysis transition state.

Indeed, the CCT2 subunit in the S2 map appeared less dynamic and better resolved than that in the NPP-state S1 map (Figs. 1d, f and Fig. 2a), indicating CCT2 may have been stabilized after ATP binding. Accordingly, the substrate density in the S2 map appeared larger than that in the S1 map, with this difference attributed to ATP-binding-induced stabilization of TRiC (Fig. 1h and Fig. 2j). Notably, in TRiC-tubulin-S2, the substrate density was found to be closely associated with all the three domains of CCT6/8 and to form loose contacts with the E domains of the other subunits (Fig. 2j) as well as with the central tail region (Fig. 2b). Compared with TRiC-tubulin-S1, it appears that ATP-binding could drive tubulin to translocate from the E domains of all the subunits slightly up towards the A/I domains, to converge more on the CCT6/8 subunits, while nevertheless remaining bound to all of the E domains (Fig. 2j).

### ATP-driven TRiC ring closure is the determinant step for tubulin folding

Importantly, inspection of our TRiC-tubulin-S3 map revealed a well resolved tubulin density in one chamber of TRiC hanging underneath its dome predominantly on the CCT6 hemisphere subunits (including CCT1/3/6/8)^21, 28^ (Fig. 2d-e and Fig. 3a-b). The tubulin density appeared to resemble the conformation of native tubulin (Fig. 3b), and hence we defined the captured tubulin as being in the near-natively folded state. Overall, the N and C domains of tubulin were relatively well resolved with atomic details observed (Fig. 3c). While the majority of the I domain of tubulin was captured (Fig. 3b), a small portion (sequence: 242-249, 279-283, 319-348) facing the central chamber was less well resolved (Fig. 3a, d), implying an intrinsically dynamic nature for this region. Interestingly, this dynamic region was noted to overlap with its interaction interface with α-tubulin to form a tubulin heterodimer or with cofactor A for the release of tubulin^43, 52^ (Fig. 3d), indicating this intrinsically dynamic region of the tubulin I domain may only be stabilized by engaging with these factors. Substantiating this speculation, our XL-MS analysis showed that tubulin I domain Lys252 forms crosslinks with all subunits of TRiC with the cross-linked Cα-Cα distances ranging from 20 to 61 Å (Fig. 3e), indicating a highly dynamic structure in this region. These results were also in line with a recent report showing reovirus σ3 capsid protein acting as a dynamic substrate within the TRiC chamber^28^. Note that this Lys252 had been shown to play an essential role in interacting with the γ-phosphate of the GTP in the α-tubulin portion of the tubulin dimer^43^ (Fig. 3f). Collectively, our data suggested that the I domain region of β-tubulin is intrinsically dynamic since it is “born” before associating with α-tubulin to form the tubulin dimer.

**Fig. 3.**
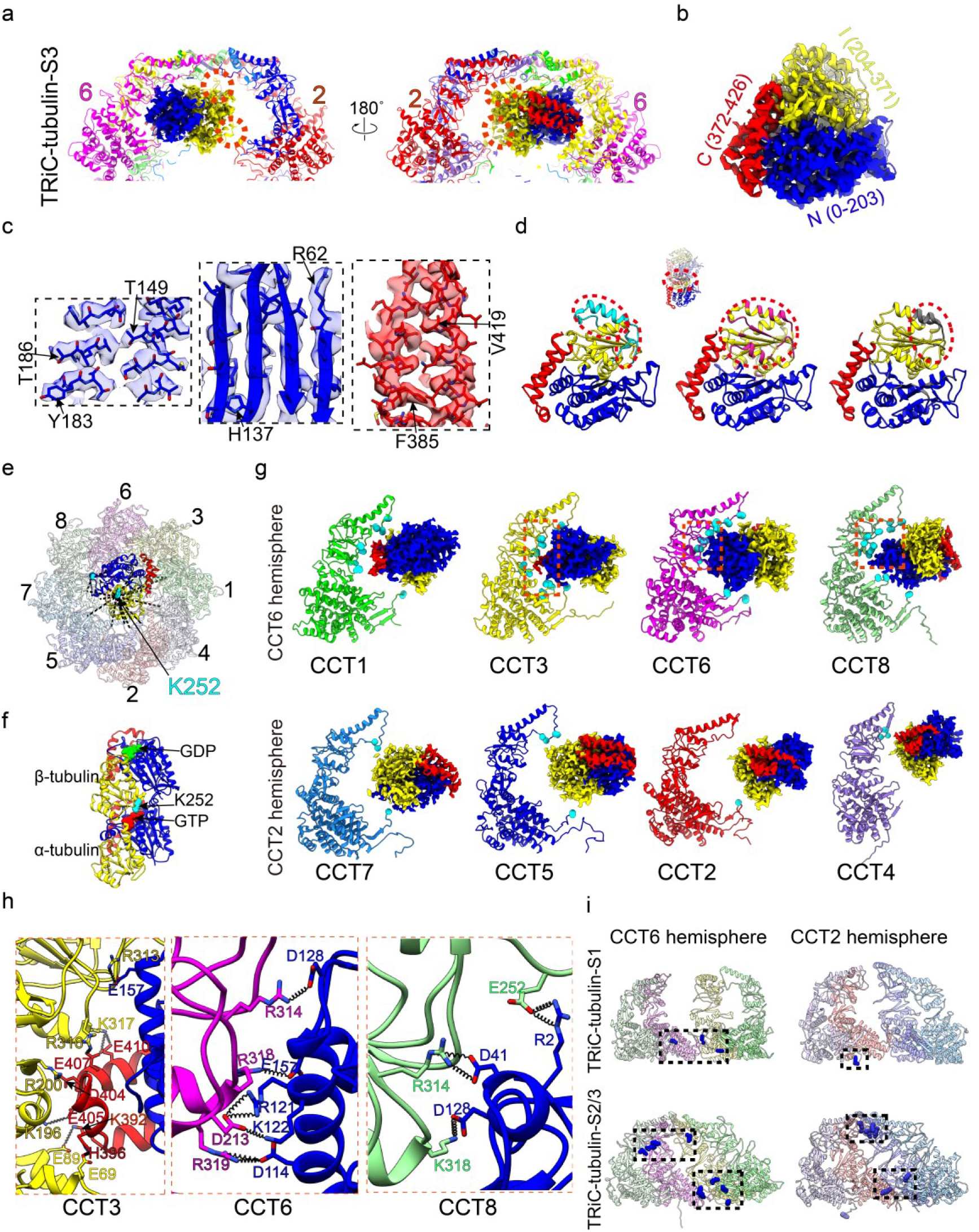
TRiC-tubulin-S3 structure showing a near-natively folded tubulin attached to the cis-ring of TRiC. **a**, Enlarged views of the engaged tubulin within the TRiC chamber in the S3 state. The tubulin N/I/C domains are rendered in blue, yellow and red, respectively, which color scheme was followed throughout. Red dotted circles indicate dynamic regions of the I domain of tubulin. **b-c**, The resolved β-tubulin inside the cis-ring of TRiC-tubulin-S3 (**b**), and its high-resolution structural features (**c**). **d**, Ribbon diagram depictions showing the unresolved dynamic portion of the tubulin I domain (in cyan, left) overlapping with its interaction interface with α-tubulin (in violet red, a distance cutoff of 4 Å, middle) or the interaction site with cofactor A (in gray, right)^52^. **e**, Mapping of the detected XLs made by tubulin K58 and K252 (cyan spheres) with the TRiC subunits in the cis-ring of TRiC-tubulin-S3. Note that every TRiC subunit made at least one such XL. **f**, Ribbon diagram illustrating the αβ-tubulin dimer structure, with β-tubulin facing upwards. K252 in β-tubulin and bound nucleotides are shown as spheres. **g**, Ribbon diagram depictions of the association between each TRiC subunit and tubulin in S3, showing the close associations of the N/C domains of tubulin with CCT6 hemisphere subunits CCT1/3/6/8, and loose associations of the tubulin I domain with CCT2 hemisphere subunits CCT7/5/2/4. Cα atoms of the TRiC amino acid residues within 4 Å distance with tubulin are shown as cyan balls. **h**, Magnified views of the regions indicated with red dotted frames in **g** to show the salt bridge interactions formed between tubulin and the CCT3/6/8 subunits. **i**, XL-MS-analysis-derived sites on TRiC (blue spheres) crosslinked with tubulin and mapped onto the corresponding indicated TRiC structure. This analysis suggested a shift in the interaction locations induced by ATP binding/hydrolysis.

Inspection of the S3 map showed that tubulin engages with TRiC mainly through its N/C domains, forming intimate salt-bridge-and H-bond-mediated contacts with all three domains of the CCT6 hemisphere subunits (Fig. 3g, h and Supplementary Table 4). The tubulin I domain also loosely interacts with CCT2 hemisphere subunits through association with the A-domain protrusion loop and C-termini of CCT7/5/2/4 (Fig. 3g). Indeed, tubulin showed an obviously larger interaction area with the CCT6 hemisphere subunits than with the CCT2 hemisphere ones (Extended Data Fig. 7l). Furthermore, inspection of S3 structure also unprecedently revealed the protrusion loop, loop^H^^10^, loop^H9^, C-terminus, and stem loop to be the main structural elements of TRiC involved in its interaction with tubulin (Fig. 3g-h and Extended Data Fig. 7m-n). Our TRiC-tubulin-S3 structure appeared overall comparable with a recent structure of closed TRiC-tubulin complex^53^, in terms of tubulin orientation and binding location within TRiC chamber.

In addition, our XL-MS data suggested that before adding nucleotide, tubulin only crossed linked with the E domains of TRiC; while in the presence of ATP-AlFx, additional XLs formed with the A/I domains of TRiC (Fig. 3i), indicating an upward shift of tubulin induced by ATP binding and hydrolysis. Indeed, more TRiC A and I domain regions were observed to be involved in the interaction with tubulin in the S2 and S3 states (Figs. 2j and 3g) than in the S1 state (Fig. 1h). Inspection of the S2 and S3 maps suggested that ATP-hydrolysis could induce considerable downward rotation of the A and E domains of TRiC from the open to closed state (Extended Data Fig. 7o). These substantial movements could dramatically reduce the chamber volume, and hence restrain the conformational landscape of the substrate and lead to a more stabilized substrate through intimate interactions with the TRiC A/I domains. In the meanwhile, the mechanical force generated from TRiC ring closure could provide extra energy to help the substrate overcome the energy barrier and transform towards the global energy minimum. This may be a general mechanism for TRiC-mediated substrate folding.

### Tubulin engages with closed TRiC through both electrostatic and hydrophilic interactions

We then inspected the closed TRiC-tubulin-S3 structure to derive the interaction properties between tubulin and TRiC. Here the electrostatic surface property inside the TRiC chamber was observed to be asymmetric, with the CCT6 hemisphere more positively charged and the CCT2 hemisphere relatively negatively charged (Fig. 4a-b), in line with previous reports on yeast and bovine TRiC^21^. Interestingly, the engaged tubulin N/C domains appear negatively charged, complementary with the contacting CCT6 hemisphere (Fig. 4b). This observation was also consistent with our findings that numerous salt bridges formed between TRiC CCT6 hemisphere subunits and tubulin N/C domains (Fig. 3h). The electrostatic surface of the tubulin I domain was found to be not complementary with the related region of TRiC (Fig. 4a-b), consistent with the lack of a strong interaction between them (Fig. 3g, Extended Data Fig. 7l). Taken together, our S3 structure revealed previously unreported strong electrostatic interactions between the tubulin N/C domains and the TRiC CCT6 hemisphere subunits.

**Fig. 4.**
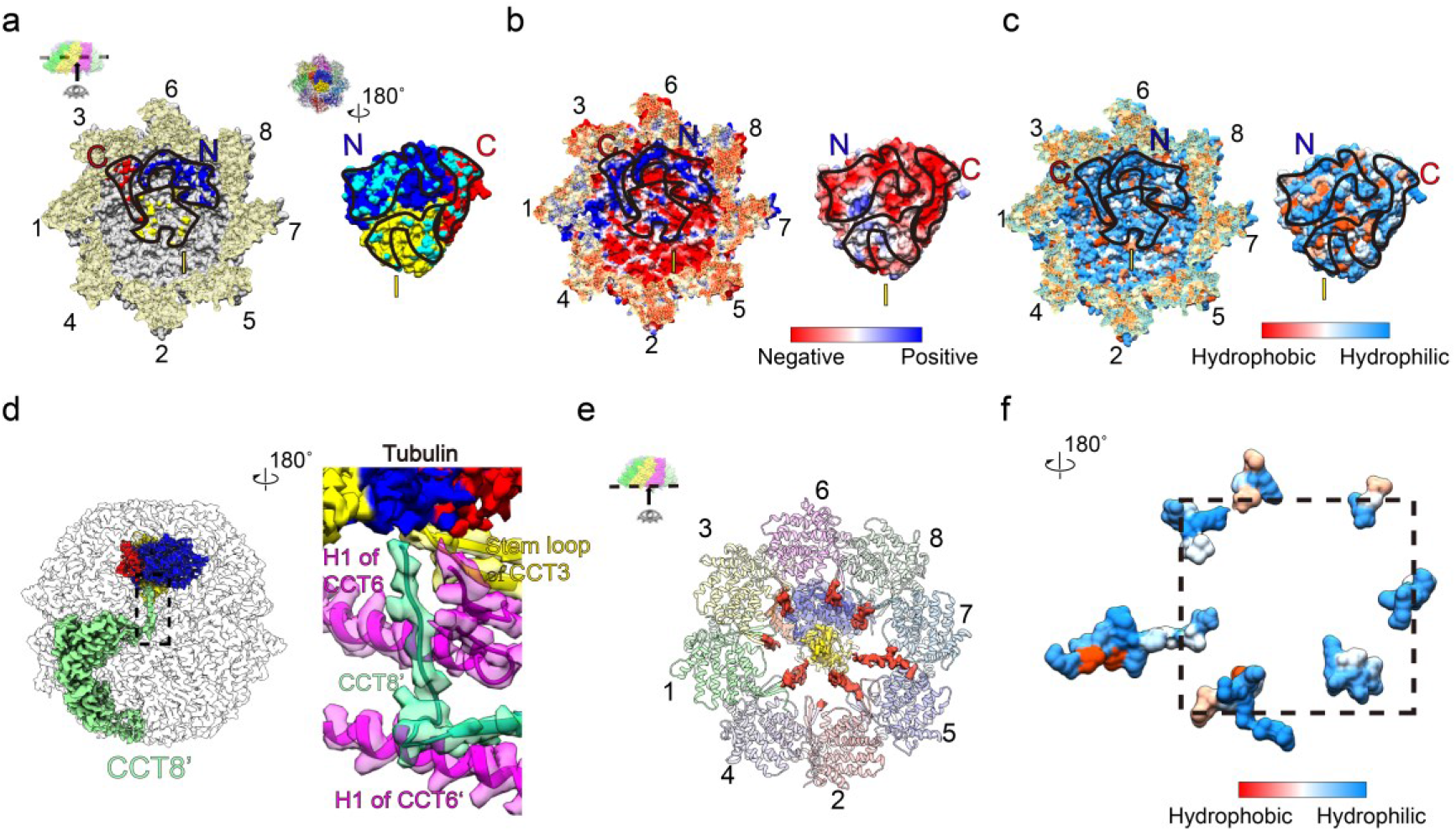
Engagement of tubulin with TRiC through electrostatic and hydrophilic interactions, and the roles of TRiC termini in tubulin folding in TRiC-tubulin-S3. **a**, Interaction interface between TRiC and tubulin. The visualization direction and region are shown in the inset. The contact surface residues of TRiC within 4 Å distance of the N/C/I domains of tubulin are colored in blue, red, and yellow, respectively (left panel, black outlines indicate the tubulin footprint on the TRiC interior cavity wall). The contact surface residues of tubulin in proximity to TRiC are colored in cyan (right panel, black outlines indicate the TRiC footprint on the tubulin structure). **b**, An electrostatic surface property analysis, suggesting complementary electrostatic interaction between the CCT3/6/8 subunits of TRiC (mainly positively charged) and the N/C domains of tubulin (mostly negatively charged). **c**, The hydrophilicity/hydrophobicity analysis between TRiC and tubulin, suggesting a hydrophilic interaction between them. **d**, Magnified view of the interaction network of the CCT8’ N-terminus. **e**, Depiction showing those resolved portion of C-termini (red density) for all TRiC subunits (besides CCT4) were observed to form contacts with tubulin (central transparent density). **f**, Depiction of the hydrophilicity/hydrophobicity of the resolved C-termini of TRiC, showing these termini to be enriched in hydrophilic residues.

Inspection of the S3 structure also suggested an enrichment of hydrophilic residues at the interaction interfaces between the TRiC inner dome and tubulin, creating a mostly polar TRiC-tubulin interface in the closed chamber (Fig. 4c). This type of polar interface was also seen in the TRiC-σ3 complex^30^. Taken together, our data suggested that the closed-state TRiC interacts with its obligate substrate tubulin through a combination of electrostatic and hydrophilic interactions to stabilize this substrate and facilitate its folding.

We have previously showed that yeast TRiC can close both rings in the presence of natural nucleotide ATP^16^. Here we performed further cryo-EM study on human TRiC incubated with ATP at 1 mM, a physiological concentration of ATP in the cell^16, 54^, and obtained two maps, including an open-state TRiC and a both-ring-closed TRiC (27.2 % of the population) (Extended Data Fig. 8a-c). The closed TRiC map showed that all nucleotide pockets were occupied, in addition to a tubulin density within one TRiC chamber in a position and orientation similar to that observed in the TRiC-tubulin-S3 map (Extended Data Fig. 8d-e). Collectively, these results suggested that the double-closed state for both yeast and human TRiC indeed exists in the presence of natural nucleotide, in line with a recent *in situ* cryo-ET study on neuron cells^55^.

### Cryo-EM map of TRiC-ADP

To capture the complete TRiC ATPase cycle, we determined a cryo-EM map of TRiC in the presence of ADP to a resolution of 3.3 Å (termed TRiC-ADP, Fig. 5a-b, Extended Data Fig. 1e, 9, and 10a-b). In this map, TRiC is in the open conformation, overall similar to that of TRiC-NPP (Extended Data Fig. 10c), but with all of the subunits loaded with ADP (Fig. 5c, Extended Data Fig. 10d). However, other than the remaining unstructured tail density, there was no engaged tubulin anymore within TRiC chamber (Fig. 5b). Corroborate to this, our XL-MS data showed no detected cross-links between TRiC and tubulin after the ADP incubation (Fig. 5d), indicating that under our experimental condition ADP binding on TRiC could potentially contribute to a release of tubulin from TRiC regardless of tubulin folding status. We performed further 3DVA on the dataset and found TRiC-ADP to be very dynamic, with all the subunits (including the most stable CCT6) displaying outward/inward tilting motions (Extended Data Fig. 10e, Supplementary Video 2). We then postulate that the dynamic nature of TRiC-ADP may contribute to the substrate release in this state. Still, we should mention that here we used the TRiC-ADP structure to mimic the state that, after releasing γ-phosphate, TRiC reached the open state and then also released the substrate.

**Fig. 5.**
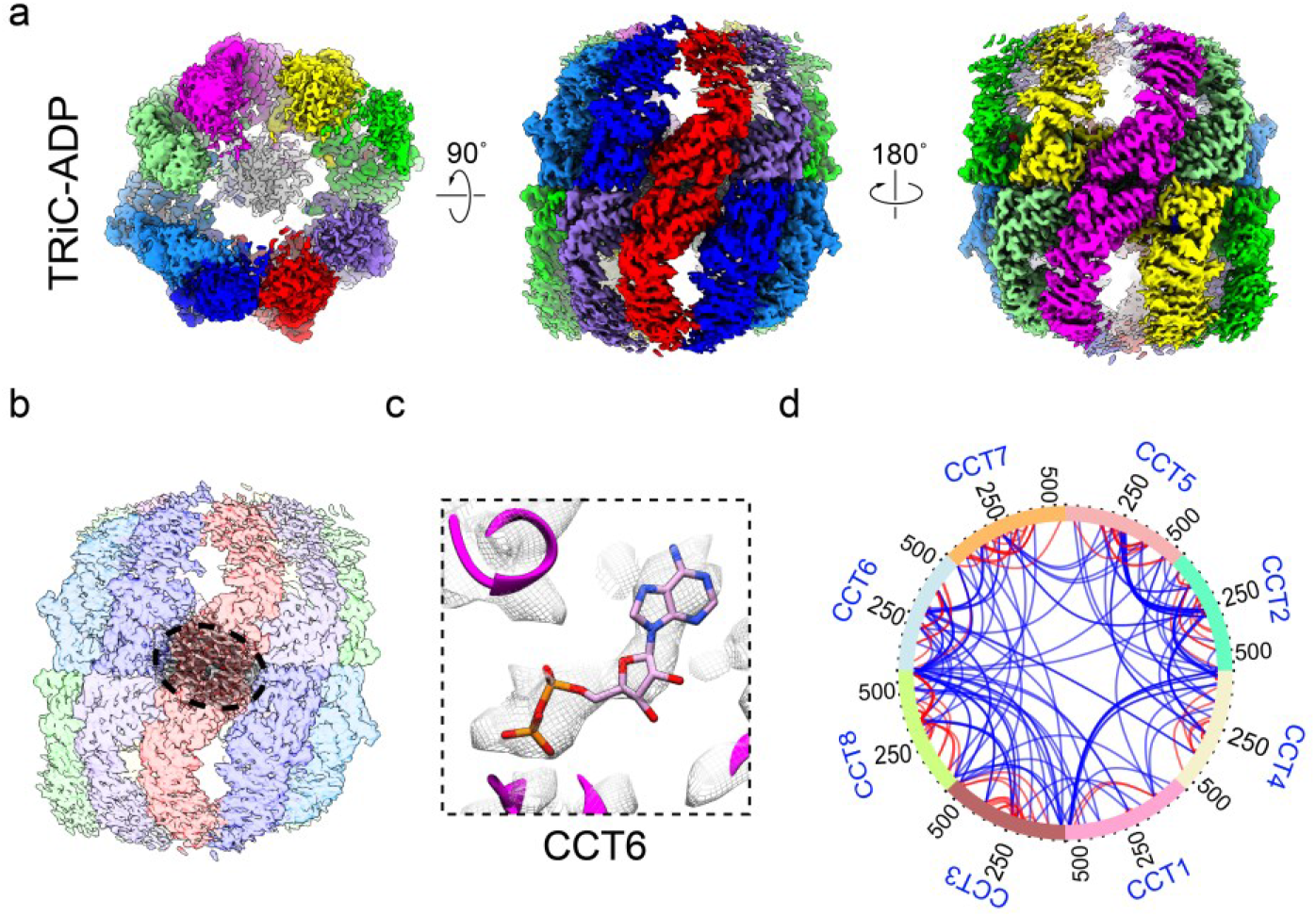
Cryo-EM and XL-MS analyses of TRiC in the presence of ADP. **a-b**, Cryo-EM map of TRiC-ADP (**a**), showing the tail density (in grey) remained but substrate released (**b**). **c**, Representative nucleotide density (in the ADP state) of CCT6 in TRiC-ADP. **d**, The XL-MS data showing no cross-links were detected between TRiC and tubulin after ADP incubation. Identified intra-subunit XLs are in red, and inter-subunit XLs in blue.

### The C- and N-termini of TRiC play distinct roles in substrate folding and TRiC allosteric coordination

Here, we showed that in each of the four open-state TRiC structures (TRiC-NPP, S1, S2, and TRiC-ADP), there is a central unstructured tail density between the two equators (Fig. 1e, g, Fig. 2b and Fig. 5b). Moreover, in both the S1 and S2 structures, the resolved portions of the TRiC C-termini and the extended unstructured tail density apparently form contacts with the tubulin substrate and hence potentially stabilize the substrate (Fig. 1g-h and Fig. 2b, j).

Strikingly, in the closed TRiC-tubulin-S3 structure, the N-terminus of almost all of the TRiC subunits were stabilized and very well resolved (Extended Data Fig. 11). Specifically, the N-termini of all subunits except CCT4 were observed to contact H1 of the neighboring subunit, and those of CCT2/4/5/7/8 to involve in the inter-ring allosteric network (Extended Data Fig. 11a-g). This observation is consistent with our previous finding in yeast TRiC, suggesting that the extra layer of N-terminal allosteric network could play important roles in TRiC ring closure^16^. Intriguingly, we observed the N-terminus of CCT8’ from the trans-ring adopting a bent conformation and extending all the way to the cis-ring to associate with the tubulin N domain (Fig. 4d), in addition to being involved in the inter-ring allosteric network by linking the CCT6/6’ N-termini and CCT3 stem loop together. This indicated that CCT8 may play a critical role in substrate stabilization/folding besides being involved in TRiC inter-ring cooperativity.

Notably, in the closed S3 state, we also resolved a portion of the C-terminal extensions for most TRiC subunits (except CCT4), and captured the uncharacterized direct interactions between these C-terminal extensions of TRiC and tubulin (Fig. 4e). These C-terminal extensions, mostly hydrophilic (Fig. 4f), stretch out from the surrounding subunits towards the center of TRiC chamber to form physical contacts with tubulin, and hence appearing like a net holding and stabilizing tubulin within the TRiC chamber (Fig. 4e), which may facilitate tubulin folding. Consistent with this, previous studies suggested that the C-termini of GroEL form contacts with its substrate, which could enhance and accelerate substrate folding, and that the hydrophilic residues on the C-termini are critical for substrate folding^56–59^.

## Discussion

Tubulin is the building block of the microtubule, which is critical to many cellular processes, and the eukaryotic chaperonin TRiC is required for tubulin biogenesis. Here, we captured snapshots of tubulin folding pathway mediated by TRiC, by acquiring six cryo-EM structures of human TRiC in its ATPase cycle with three of them engaged with endogenous tubulin in different folding stages (Fig. 1, 2, and 5). Importantly, our TRiC-tubulin-S3 structure revealed a near-natively folded tubulin engaged with closed TRiC in one chamber, primarily with the A/I domains of the CCT3/6/8 subunits through electrostatic and hydrophilic interactions (Fig. 4a-c). We also noticed the interaction of TRiC termini with tubulin in the open TRiC-tubulin-S1 and -S2 states (Fig. 1h, 2j), and found that in the closed S3 state the N terminus of CCT8 from the opposite ring and the C-terminal extensions of almost all TRiC subunits may play a role in tubulin stabilization and folding (Fig. 4d-f).

### Proposed pathway and mechanism of TRiC-mediated tubulin folding

Here we propose a complete picture of TRiC-mediated tubulin folding pathway and mechanism conjugated with the ATP-driven conformational cycle of TRiC (Fig. 6a). After being translated from ribosomes, nascent tubulin polypeptides are delivered to TRiC by co-chaperon prefoldin to enter its folding pathway associated with the TRiC ATPase cycle^49, 60^. Subsequently, tubulin can be released inside TRiC chamber, making contacts with the E domains of all of the TRiC subunits and with the unstructured termini of TRiC, to form the TRiC-tubulin-S1 state, in which the tubulin is relatively dynamic (step 1). After ATP binds to TRiC, with the resulting stabilization of the folding machinery, tubulin can be gradually translocated upwards to associate with the A/I domains of CCT6/8 in addition to keeping contacts with the E domains of all of the subunits (step 2). Once hydrolysis of ATP triggers TRiC ring closure (with the involvement of TRiC N-termini), C-terminal extensions of TRiC and the N-terminus of CCT8 from the opposite ring could stabilize and potentially facilitate the upward translocation of tubulin to result in the formation of intimate electrostatic and hydrophilic interactions between the A domain of CCT6 hemisphere subunits and the N/C domains of tubulin. These directional contacts and constraints on the tubulin N/C domains could facilitate its folding towards the native state, and the mechanical force generated by TRiC ring closure including the considerable downward rotation of the A domain of every TRiC subunit could be propagated to tubulin and drive it to overcome the energy barrier to transform towards the global minimum reaching the folded state (step 3). Subsequently, in the γ-phosphate released open TRiC-ADP state, the associated tubulin could be released from TRiC chamber (step 4). Based on previous reports, the dynamic β-tubulin I domain could then be recognized and capped by cofactor A, followed by the assembly of β-tubulin with α-tubulin into tubulin heterodimers with the assistance of cofactors C, D, and E^61–63^. Finally, the tubulin heterodimers assemble into the microtubule, which goes on to perform its biological functions.

**Fig. 6.**
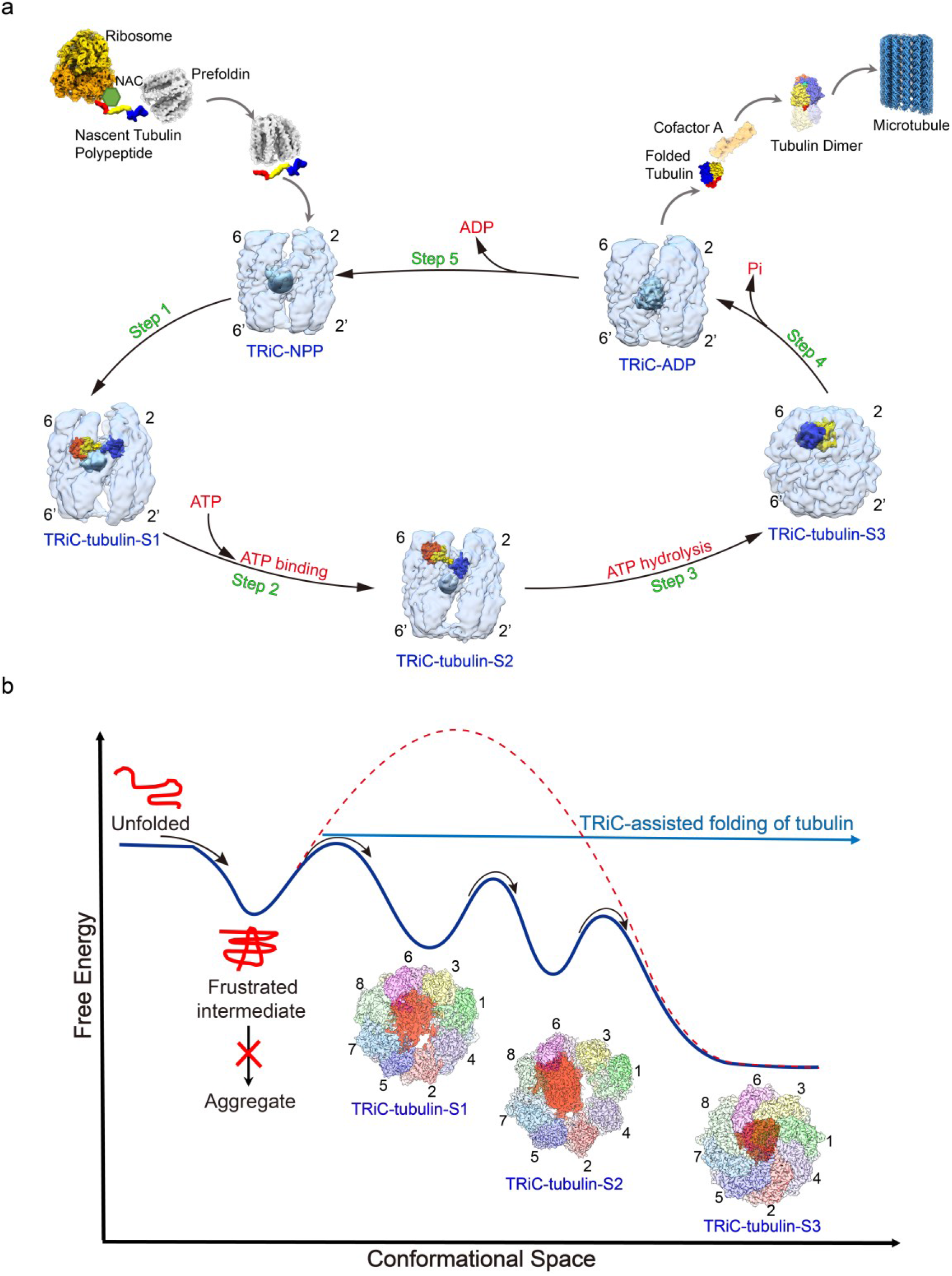
The proposed pathway and mechanism of TRiC-mediated tubulin folding. **a**, The proposed pathway of tubulin folding mediated by TRiC associating with its ATP-driven conformational cycle. After being translated from ribosome, nascent tubulin polypeptides are delivered to TRiC-NPP by co-chaperon prefoldin. TRiC is shown in transparent light blue, with its unstructured termini in sky blue density and CCT2/6 being labeled. Tubulin is then released inside the TRiC chamber, contacting with the E domains of all TRiC subunits and with the unstructured termini of TRiC, to form the TRiC-tubulin-S1 state (step 1). After ATP binding to TRiC, tubulin can gradually translocate upwards to associate with the A/I domains of CCT6/8 in addition to keeping contacts with the E domains, forming the TRiC-tubulin-S2 state (step 2). Once ATP-hydrolysis triggers TRiC ring closure, forming the TRiC-tubulin-S3 state, the generated mechanical force together with the directional contacts and constraints on tubulin could facilitate tubulin folding towards the native state (step 3). Subsequently, in the γ-phosphate released open TRiC-ADP state, the associated tubulin could be released from TRiC chamber (step 4). The dynamic β-tubulin I domain could be capped by cofactor A, then assembled with α-tubulin into tubulin heterodimers. Finally, the tubulin heterodimers assemble into the microtubule. TRiC could then release ADP to resume to the NPP state (step 5). All TRiC maps were low passed to 8 Å for easier visualization. **b**, Hypothetical energy landscape of tubulin folding assisted by TRiC. Without TRiC assistance, nascent tubulin is prone to form aggregates or needs to overcome a high energy barrier to achieve its native state. Engagement of tubulin with TRiC in a manner associated with the TRiC ATPase cycle could potentially confine the energy landscape of tubulin and lower its energy barrier, resulting in stabilization of tubulin and finally becoming folded into a substantially near-native state.

### Conformational landscape of TRiC-mediated tubulin folding

Without TRiC-assisted folding, nascent tubulins are prone to form aggregates due to the high energy barrier for spontaneous folding to reach the native folded state^64^. In the current work, by capturing multiple intermediate states of tubulin folding assisted by TRiC, we were able to derive a hypothetical energy landscape of the three distinct states of TRiC-tubulin on the basis of nucleotide binding status, complex stability, and population distribution (Fig. 6b). In TRiC-tubulin-S1, the engagement with TRiC could potentially stabilize the nascent tubulin, leading it to overcome the first energy barrier to reach a lower level in the energy landscape. Still, the tubulin at this stage remains quite dynamic, accompanying the dynamic nature of TRiC in the NPP state with only CCT3/6/8 subunits preloaded with ADP (Extended Data Fig. 3f-g). The subsequent binding of ATP to all of the TRiC subunits could drive TRiC and associated tubulin to overcome the next barrier and to stabilize in the steadier TRiC-tubulin-S2 state, evidenced by the observation that the engaged tubulin density here becoming further stabilized and appearing larger. Only when ATP-hydrolysis provides sufficient chemical energy to enable TRiC to overcome the final energy barrier, shutting both rings and becoming transformed to the most stable TRiC-tubulin-S3 state, the confined tubulin is now folded to its near native state inside the closed TRiC chamber. This step may be the rate-limiting step in tubulin folding process. Thus, the ATP-mediated gradual stabilization of TRiC guides tubulin along a pathway that avoids deep kinetic traps in the folding energy landscape. That is to say, binding of tubulin to TRiC could lower the folding energy barrier of tubulin and result in its stabilization, so that it eventually reaches its folded state accompanying ATP-driven TRiC ring closure.

### Asymmetric working mechanism of TRiC

TRiC subunits have been previously demonstrated to display a gradient of ATP affinities^27^, as well as differences in nucleotide binding^15, 19, 28^ and consumption^16^, with these features closely related to the structural asymmetry among TRiC subunits^19, 24, 65^. In the current study, our results further depicted that in the TRiC-NPP state, the bound ADP in CCT3/6/8 may play a role in stabilizing these subunits, making them appear less intrinsically dynamic than the other subunits (Fig. 1d, f). Consequently, CCT3/6/8 may serve as a dock for an initial engagement of tubulin (Fig. 1h and Fig. 2j), and eventually for close electrostatic and hydrophilic contacts with the N/C domains of tubulin in its nearly folded state (Fig. 3h and Fig. 4a-c). Consistently, previous studies also suggested important roles of CCT3/6/8 in the recognition of other substrates such as mLST8, reovirus σ3 capsid protein, and AML1-175^26–28^. Taken together, the asymmetry of TRiC subunits in structural features and nucleotide consumption may contribute to TRiC-assisted substrate folding.

In addition, we found that along with the substrate folding, from a relatively disordered state to an ordered state, TRiC also appears to follow a similar trend: transforming from the relatively dynamic and asymmetric open S1 state to the stabilized, rather symmetrical both-ring-closed S3 state. And during this process, the nucleotide occupancy status also appears to change, from an initial partial occupancy in the NPP state to the full occupancy in all the subunits of TRiC, with the nucleotide states becoming more homologous (Extended Data Fig. 3f, 7g-h). In the nucleotide binding and consumption, as well as ring closure, the asymmetric machinery—TRiC displays a stepwise mechanism^16, 19^ rather than a concerted manner as in the Group I system^66, 67^. As has been suggested previously^68^, the asymmetry or non-concerted nature of TRiC in the open state could be beneficial for its ability to recognize and engage with diverse substrates. Moreover, despite that our study and most other studies revealed only one substrate engaged with TRiC^7, 11, 28, 29, 69, 70^, there was a crystal structure showing that TRiC can simultaneously bind two tubulins, one per ring^15^, implying two substrates could interact with a TRiC complex simultaneously at least under certain physiological conditions^68^ and both rings of TRiC may have the ability to perform substrate folding concurrently for efficient protein quality control under these conditions.

In summary, our study depicted a thorough picture of the pathway and conformational landscape of TRiC-mediated tubulin folding accompanying the ATPase cycle of the folding machinery. Furthermore, our determination of the interaction sites between tubulin and the closed TRiC chamber is beneficial for the development of novel and effective therapeutic agents specifically targeting TRiC-tubulin interactions.

## Methods

### Purification of human TRiC

Human TRiC was purified from HEK293F cells according to the published protocol with some modifications^28, 49, 71^. Briefly, the pellet was lysed with iced MQA-10% glycerol buffer (50 mM NaCl, 20 mM Hepes pH 7.4, 5 mM MgCl2, 1 mM DTT, 10% glycerol, 1 mM PMSF, and 2 mM ATP) using one Protease Inhibitor Cocktail Tablet (Roche) per 100 mL of the lysate. The lysed material was then subjected to centrifugation (20,000 g for 30 min to remove cellular debris and nuclei and then 140,000 g for 1.5 h to remove the ribosome) in order to separate the cytoplasmic fraction. The resulting supernatant was filtered using a 0.44 μm filter membrane, and then passed through a Q Sepharose column (GE Healthcare). TRiC was eluted in a gradient from 40 to 80% MQB-5% glycerol (MQA with 1 M NaCl). The fractions containing TRiC were pooled, diluted with MQA to ensure an NaCl concentration of about 100 mM, and applied to a Heparin HiTrap HP column (GE Healthcare). TRiC was eluted in a gradient from 20 to 65% MQB-5% glycerol. The fractions containing TRiC were pooled and incubated with 10 mM ATP on a shaker (220 rpm, 37 °C, 30 min) to allow TRiC to cycle and release substrate before performing gel filtration chromatography (GFC). The sample was then concentrated down to 0.5 mL and loaded onto a Superose 6 Increase 10/300 GL column (GE Healthcare) with MQA-5% glycerol without ATP. TRiC eluted at about 13-15.5 mL of the size-exclusion column, consistent with that of a 1-MDa complex. Finally, the sample was subjected multiple times to buffer exchange to remove remaining ATP in the buffer, and we obtained biologically active TRiC (Extended Data Fig. 1a-b).

It is noteworthy that TRiC-mediated folding of tubulin is associated with release of predominantly nonnative forms of tubulin from chaperonin, and with the majority of released tubulin requiring further rounds of binding/release to reach its native state^72^. This indicates the released nonnative substrate could re-bind on TRiC to go through further round of ATPase cycle. In our experimental condition, it is possible that the tubulin in TRiC-tubulin-S1 has already gone through one or several ATPase cycles.

### ATPase activity assay

The ATP hydrolysis rate of TRiC was measured by performing an NADH-coupled assay^73^. In this assay, each ATP hydrolysis event allows for a conversion of one molecule of phosphoenolpyruvate into pyruvate catalyzed by pyruvate kinase, with pyruvate then converted to lactate by L-lactate dehydrogenase, which results in oxidation of a single NADH molecule. Loss of NADH over time, a measure quantifiably proportional to the ATP hydrolysis rate, was monitored in the current work by measuring the decrease in absorbance of light at a wavelength of 340 nm. All of the assays were conducted at room temperature in a buffer containing 10 mM HEPES/NaOH pH 7.5, 50 mM NaCl, and 10 mM MgCl2, in the presence of 1 mM ATP. Experiments were performed in triplicate using 0.3 μM of the protein complex. Absorbance was measured in a 200 µl reaction volume using a 96-well plate reader. Data analysis was performed using Graph Pad Prism 8.

### ADP/ATP ratio assay

To identify the form of the nucleotide in our purified human TRiC, we carried out luciferin-luciferase reactions with an ADP/ATP ratio assay kit^74^ (Sigma-Aldrich). To release the bound nucleotide from TRiC for measurement, the TRiC sample was first digested with proteinase K, according to a previously published protocol^75^ with minor modifications. TRiC proteolytic-digestion experiments were carried out by adding a 1 µl aliquot of 2 mg/ml proteinase K to a 19 µl aliquot of 4.5 mg/ml TRiC in dilution buffer (20 mM Hepes-KOH pH 7.4, 50 mM NaCl, 5 mM MgCl2, 1 mM DTT, and 5% glycerol). The digestion was performed at 37 °C for 1 h. The reactions were then terminated by adding PMSF (to a final concentration of 5 mM) into the reaction mixture and waiting for 10 minutes. Afterwards, the nucleotide form in the TRiC sample was identified by using the ADP/ATP ratio assay kit according to the manufacturer’s protocol, and the RLU values of ATP and ADP were measured with a Synergy Neo2 multimode reader (BioTek).

### Cross-linking and mass spectrometry analysis

The purified TRiC was cross-linked by using bis[sulfosuccinimidyl] suberate (BS^3^) (Sigma), with a spacer arm of 11.4 Å between their Cα carbons, on ice for 2 hours. The final concentration of the crosslinker was 2 mM. The reaction was then terminated by using 50 mM Tris-HCl pH 7.5 at room temperature for 15 minutes. For the sample containing ATP-AlFx, the purified TRiC was incubated with 1 mM ATP, 5 mM MgCl2, 5 mM Al(NO3)3, and 30 mM NaF for 1 h at 37 °C, and the resulting product was cross-linked by using BS^3^ following the above-mentioned conditions. Cross-linked complexes were precipitated and digested for 16 hours at 37 °C by using trypsin at an enzyme-to-substrate ratio of 1:50 (w/w). The tryptic-digested peptides were desalted and loaded on an in-house packed capillary reverse-phase C18 column (length of 40 cm, 100 µm ID x 360 µm OD, 1.9 µm particle size, pore diameter of 120 Å) connected to an Easy LC 1200 system. The samples were analyzed with a 120 min-HPLC gradient of 6% to 35% of buffer B (buffer A: 0.1% formic acid in Water; buffer B: 0.1% formic acid in 80% acetonitrile) at 300 nL/minute. The eluted peptides were ionized and directly introduced into a Q-Exactive mass spectrometer using a nano-spray source. Survey full-scan MS spectra (from m/z 300–1800) were acquired using an Orbitrap analyzer with a resolution r=70,000 at an m/z of 400. Cross-linked peptides were identified and evaluated using pLink2 software^76^.

### Cryo-EM sample preparation

To prepare the vitrified sample of TRiC, the purified TRiC was diluted to 1.2 mg/ml, and an aliquot of 2 μl of this sample was applied onto a plasma cleaned holey carbon grid (Quantifoil, R1.2/1.3, 200 mesh). The grid was blotted with Vitrobot Mark IV (Thermo Fisher Scientific) and then plunged into liquid ethane cooled by liquid nitrogen. To prepare the sample of TRiC in the presence of 1 mM ATP-AlFx, the purified TRiC was diluted to 1.2 mg/ml, and incubated with 1 mM ATP, 5 mM MgCl2, 5 mM Al (NO3)3, and 30 mM NaF at 37 °C for 1 h prior to freezing. To prepare the sample of TRiC with 1 mM ADP, the purified TRiC was diluted to 1.2 mg/ml, and incubated with 1 mM ADP and 5 mM MgCl2 at 37 °C for 1 h prior to freezing. Moreover, to prepare the sample of TRiC in the presence of 1 mM ATP, the purified TRiC was diluted to 1.2 mg/ml, and incubated with 1 mM ATP and 5 mM MgCl2 at 37 °C for 30 s before freezing. We then followed the above-mentioned procedure to prepare the vitrified sample in each case.

### Data acquisition

For each of the experimental sample conditions mentioned above except for TRiC-ATP dataset, cryo-EM movies of the sample were collected using a Titan Krios electron microscope (Thermo Fisher Scientific) operated at an accelerating voltage of 300 kV with a nominal magnification of 18,000x (yielding a pixel size of 1.318 Å, Table 1). The movies were recorded on a K2 Summit direct electron detector (Gatan) operated in the super-resolution mode under low-dose condition in an automatic manner using SerialEM^77^. The exposure time for each frame was 0.2 s and the total accumulation time was 7.6 s, leading to a total accumulated dose of 38 e^-^/Å^2^ on the specimen. For TRiC-ATP dataset, the movies were collected in a magnification of 81,000x (yielding a pixel size of 0.89 Å, Table 1) utilizing a K3 direct electron detector (Gatan) operated in the counting mode under a low-dose condition in an automatic manner using EPU (Thermo Fisher Scientific). Each frame was exposed for 0.05 s, and the total accumulation time was 2 s, leading to a total accumulated dose of 50 e^-^/Å^2^ on the specimen.

**Table 1.** Cryo-EM data collection, processing, and model validation statistics.

|  | TRiC-NPP<br>(EMDB-32922)<br>(PDB-7X0A) | TRiC-tubulin-S1<br>(EMDB-32903)<br>(PDB-7WZ3) | TRiC-tubulin-S2<br>(EMDB-32989)<br>(PDB-7X3J) | TRiC-tubulin-S3<br>(EMDB-32923)<br>(PDB-7X0S) | TRiC-ADP-AlFx<br>(EMDB-32926)<br>(PDB-7X0V) | TRiC-ADP<br>(EMDB-32993)<br>(PDB-7X3U) | TRiC-ATP-closed<br>(EMDB-33025)<br>(PDB-7X6Q) | TRiC-ATP-open<br>(EMDB-33053)<br>(PDB-7X7Y) |
| --- | --- | --- | --- | --- | --- | --- | --- | --- |
| **Data collection and processing** |  |  |  |  |  |  |  |  |
| Magnification |  |  |  | 18,000 |  |  | 81,000 |  |
| Voltage (kV) |  |  |  | 300 |  |  | 300 |  |
| Detector |  |  |  | K2 summit |  |  | K3 |  |
| Electron exposure<br>(e-/Å²) |  |  |  | 38 |  |  | 50 |  |
| Defocus range (µm) |  |  |  | -0.8 to -2.5 |  |  | -0.8 to -2.5 |  |
| Pixel size (Å) |  |  |  | 1.318 |  |  | 0.89 |  |
| Symmetry imposed |  |  |  | C1 |  |  | C1 |  |
| Initial particle images<br>(no.) | 761,665 | 761,665 | 943,101 | 943,101 | 943,101 | 884,769 | 351,262 | 351,262 |
| Final particle images<br>(no.) | 185,216 | 115,666 | 87,764 | 103,406 | 62,412 | 239,315 | 41,646 | 111,413 |
| Map resolution (Å) | 3.1 | 4.1 | 4.2 | 3.1 | 3.2 | 3.3 | 4.5 | 3.8 |
| FSC threshold | 0.143 |  |  |  |  |  |  |  |
| Map resolution<br>range (Å) | 3.0-8.0 | 3.8-8.4 | 3.8-8.4 | 2.8-4.5 | 2.9-4.5 | 3.0-8.0 | 3.5-5.5 | 3.4-6.4 |
| **Refinement** |  |  |  |  |  |  |  |  |
| Initial model used (PDB<br>code) | 5GW4;<br>5GW5 | 5GW4;<br>5GW5 | 5GW4;<br>5GW5 | 6KS6;<br>5JCO | 6KS6 | 5GW4;<br>5GW5 | 6KS6 | 5GW4;<br>5GW5 |
| Model resolution (Å) | 4.2 | 7.4 | 7.2 | 3.4 | 3.6 | 3.6 | 4.5 | 4.2 |
| FSC threshold | 0.5 |  |  |  |  |  |  |  |
| Map sharpening *B*<br>factor (Å²) | -76.7 | -133.7 | -144.4 | -53.3 | -80.4 | -96.2 | -135.8 | -111.4 |
| Model composition |  |  |  |  |  |  |  |  |
| Non-hydrogen atoms | 63,694 | 58,818 | 63,464 | 63,367 | 65,020 | 63,964 | 64,486 | 63,532 |
| Protein residues | 8,314 | 7,686 | 8,241 | 8,866 | 8,440 | 8,314 | 8,440 | 8,314 |
| Ligands | 6 | 6 | 16 | 48 | 48 | 16 |  |  |
| *B* factors (Å²) |  |  |  |  |  |  |  |  |
| Protein | 120.55 | 127.93 | 31.20 | 55.75 | 79.70 | 33.58 | 55.28 | 179.95 |
| Ligand | 57.86 | 45.81 | 11.53 | 51.16 | 72.45 | 8.06 |  |  |
| R.m.s. deviations |  |  |  |  |  |  |  |  |
| Bond lengths (Å) | 0.005 | 0.004 | 0.005 | 0.003 | 0.004 | 0.003 | 0.091 | 0.003 |
| Bond angles (°) | 0.533 | 0.996 | 1.105 | 0.665 | 1.072 | 0.655 | 1.669 | 0.886 |
| **Validation** |  |  |  |  |  |  |  |  |
| MolProbity score | 1.39 | 1.13 | 1.40 | 1.43 | 1.54 | 1.17 | 1.35 | 1.44 |
| Clashscore | 3.67 | 2.17 | 4.21 | 3.19 | 4.23 | 1.78 | 3.45 | 5.18 |
| Poor rotamers (%) | 0.04 | 0.00 | 0.07 | 0.04 | 0.04 | 0.07 | 0.01 | 0.00 |
| Ramachandran plot |  |  |  |  |  |  |  |  |
| Favored (%) | 96.50 | 97.25 | 96.82 | 95.36 | 95.10 | 96.48 | 96.68 | 97.09 |
| Allowed (%) | 3.47 | 2.75 | 3.17 | 4.64 | 4.90 | 3.43 | 3.16 | 2.91 |
| Disallowed (%) | 0.03 | 0.00 | 0.01 | 0.00 | 0.00 | 0.09 | 0.15 | 0.00 |

### Image processing and 3D reconstruction

We performed single-particle analysis mainly using Relion 3.1^78, 79^ unless otherwise specified (Table 1). All images were aligned and summed using MotionCor2^80^. After CTF parameter determination using CTFFIND4^81^, particle auto-picking, manual particle checking, and reference-free 2D classification, particles with TRiC features were remained for further processing.

For the TRiC sample before addition of nucleotides, 761,665 particles were picked from the original micrographs, and after 2D classification, 709,192 particles remained (Extended Data Fig. 2a). These particles were subjected to 3D refinement, and then were re-extracted and re-centered using the refinement coordinates. After one round of no-align 3D classification and another round of 3D classification, we then combined the substrate-containing particles (including class 1 from the first round and class 3 from the second round) together, and performed another round of 3D classification, generating a substrate-containing class with 169,098 particles. After carrying out CTF refinement and polishing on these particles, we obtained a map with weak substrate density at 4.0-Å-resolution. We subtracted the substrate density and performed a no-align 3D classification, and a good class with 68.4% of the particles were reverted to the original particles. We then performed a local refinement on these particles and reconstructed a 4.1-Å-resolution TRiC-tubulin-S1 map displaying better substrate density. For the substrate-free class 4 particles from the second round of 3D classification, we performed another round of 3D classification, resulting in a better class having 185,216 particles. After CTF refinement and Bayesian polishing, these particles were refined to a 3.1-Å-resolution TRiC-NPP map. The resolution estimation was based on the gold-standard Fourier Shell Correlation (FSC) criterion of 0.143.

For the dataset of TRiC in the presence of ATP-AlFx, after 2D classification, 544,809 particles remained (Extended Data Fig. 5a). These particles were subjected to 3D classification. The particles from class 4, displaying closed-state features, were subjected to another round of 3D classification and a resulting better class with 168,472 particles was further refined to produce a 2.9-Å-resolution map with weak substrate density inside the TRiC chamber. Thus, we focused on the extra density inside the chamber and performed a no-align 3D classification, generating 4 classes. Particles from classes 1 and 2, appearing to have tubulin density, were combined (total of 103,406 particles) and refined to produce a 3.1-Å-resolution TRiC-tubulin-S3 map; class 3 (62,412 particles), having an extra tail density, were used to reconstruct a 3.2-Å-resolution TRiC-ADP-AlFx map. In addition, particles of classes 1-3 from the first round of 3D classification were subjected to another round of 3D classification and yielded a better class with 98,046 particles that were refined to produce a 4.3-Å-resolution map. Through focused refinement and focused classification, we eventually obtained a 4.2-Å-resolution map with tubulin density in one ring, termed TRiC-tubulin-S2.

For the dataset of TRiC-ADP, after 2D classification, 344,486 and 319,306 particles remained for dataset 1 and 2, respectively (Extended Data Fig. 9a). After refinement and re-centering, the particles were further cleaned in cryoSPARC through 2D classification and heterogeneous refinement. Then the good particles were subjected to two rounds of 3D classification in Relion 3.1. Subsequently, the better-resolved classes 3/4 from dataset 1 and classes 2/3 from dataset 2 were combined and subjected to one more round of 3D classification. The better class with 239,315 particles were further refined to a 3.3-Å-resolution TRiC-ADP map.

For the TRiC-ATP dataset, after 2D classification 225,085 particles remained (Extended Data Fig. 8a). These particles were refined and re-extracted, and further cleaned up in cryoSPARC by 2D classification and heterogeneous refinement to yield three classes. Class 1 open-state particles were subjected to another round of heterogeneous refinement, and the 111,413 particles of the better class 1 were refined to produce a 3.8-Å-resolution TRiC-ATP-open map. Closed-state particles from class 2 of the first round of heterogeneous refinement were subjected to another round of heterogeneous refinement, and particles from the better class 2 were refined in Relion 3.1 to produce a 4.5-Å-resolution TRiC-ATP-closed map.

### Model building by flexible fitting

We built the homology models for human TRiC in the open and closed states and for tubulin (TUBB5) using the SWISS-MODEL server^82^, with the cryo-EM structures of the yeast TRiC in the open and closed conformations (PDB ID: 5GW4, 5GW5, 6KS6^16, 19^) and the cryo-EM structure of human TUBB3 tubulin (PDB ID: 5JCO^83^) as templates, respectively. Afterwards, we refined the model against the corresponding cryo-EM map using Rossetta^84^, and then Phenix^85^. Furthermore, to improve the fitting between model and map, we performed real-space refinement using COOT^86^. Finally, we used Phenix again for the last round of flexible fitting of the entire complex.

We used UCSF Chimera and ChimeraX for generating figures and performing electrostatic surface property calculations^87, 88^. Interaction surface analysis was conducted by using the PISA server^89^.

## Acknowledgements

We are grateful to the staffs of the NCPSS Electron Microscopy facility, Database and Computing facility, Mass Spectrometry facility, and Protein Expression and Purification facility for instrumental support and technical assistance. This work was supported by grants from the Strategic Priority Research Program of CAS (XDB37040103), the National Basic Research Program of China (2017YFA0503503), the NSFC (32130056, 31670754 and 31872714), the Shanghai Academic Research Leader (20XD1404200), Shanghai Pilot Program for Basic Research–CAS, Shanghai Branch (JCYJ-SHFY-2022-008), and the CAS Facility-based Open Research Program.

## Author contributions

Y.C. and C.L. designed the experiments. C.L. purified the proteins and performed functional analysis with the involvement of Q.Z.. C.L. and M.J. with the involvement of S.W. and W.H. collected the cryo-EM data. C.L. with the involvement of C.X. and Y-F. Wang performed data reconstruction and model building. Y-X. Wang, L.D., and L.B. helped supervise the functional analysis. C.P., C.L., and Y.Y. performed the XL-MS analysis. Y.C. and C.L. analyzed the structure and wrote the manuscript.

## Data availability

All data presented in this study are available within the figures and in the Supplementary Information. Cryo-EM maps and the associated models have been deposited in the EMDB and Protein Data Bank, respectively, with the accession ID as the following: TRiC-NPP (EMDB-32922, PDB-7X0A), TRiC-tubulin-S1 (EMDB-32903, PDB-7WZ3), TRiC-tubulin-S2 (EMDB-32989, PDB-7X3J), TRiC-tubulin-S3 (EMDB-32923, PDB-7X0S), TRiC-ADP-AlFx (EMDB-32926, PDB-7X0V), TRiC-ADP (EMDB-32993, PDB-7X3U), TRiC-ATP-closed (EMDB-33025, PDB-7X6Q), TRiC-ATP-open (EMDB-33053, PDB-7X7Y).

## Competing interests

The authors declare no competing interests.

---

**Extended Data Fig. 1.**
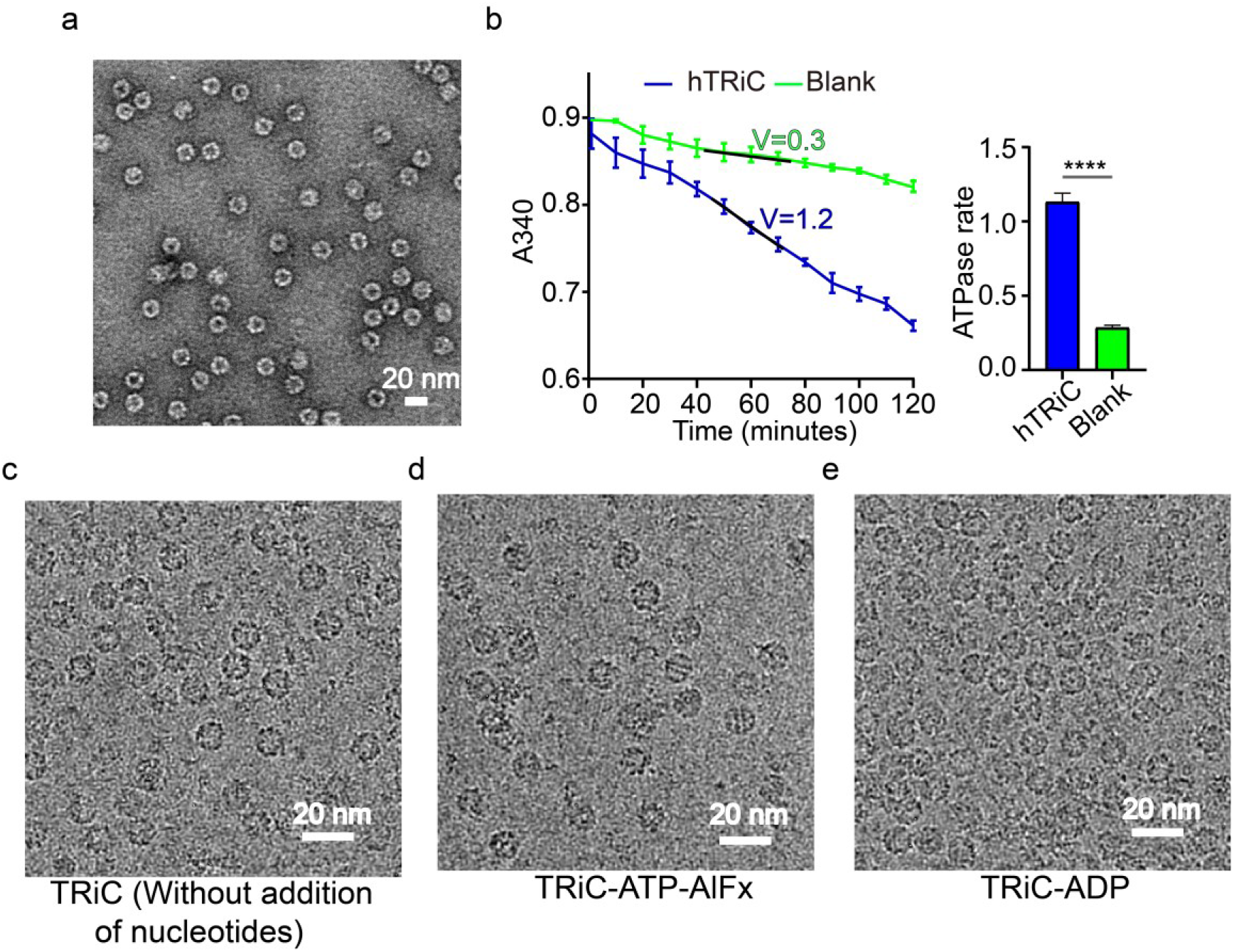
Human TRiC purified from HEK293F cells. **a**, Representative negative-stain EM image of purified TRiC. **b**, NADH-coupled enzymatic assays of our purified TRiC and a blank (TRiC buffer without proteins). The results revealed that TRiC was biologically active and could hydrolyze ATP. The ATPase rates (V, with unit of “mole ATP/ [mole TRiC • min]”) determined by fitting the linear part of the ATP hydrolysis reaction curves are also provided. An analysis of the significance of the difference between the results (right) suggested that the TRiC sample showed a significantly higher ATPase activity than did the blank, with a statistical significance of ****P < 0.0001. For all quantifications, data were plotted as mean ± SD for three independent replicates. **c-e**, Representative cryo-EM images of TRiC sample before adding nucleotides (**c**), in the presence of ATP-AlFx (**d**), and in the presence of ADP (**e**).

**Extended Data Fig. 2.**
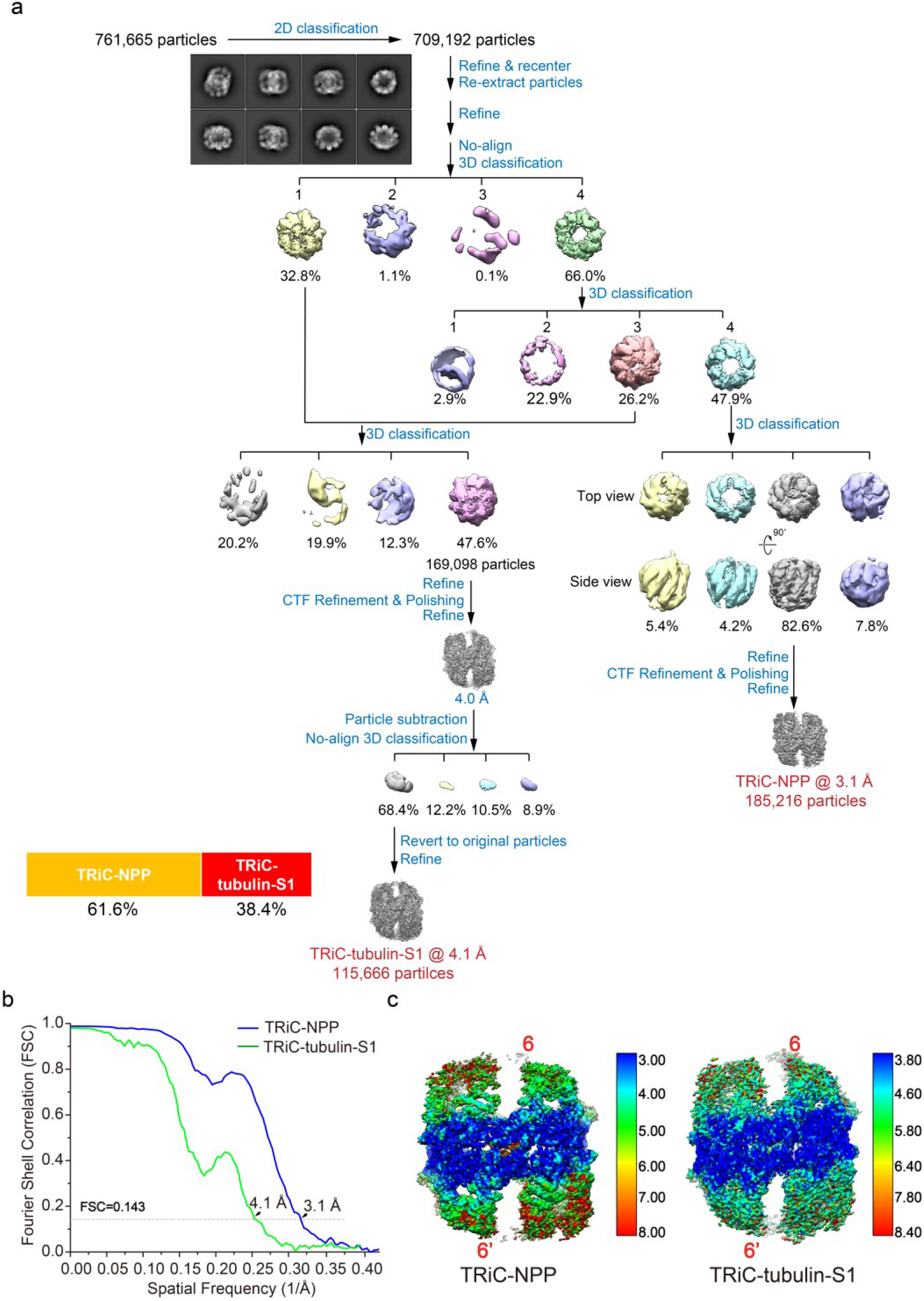
Work flow for the processing of the cryo-EM data of endogenously purified TRiC. **a**, Procedure used to process the TRiC cryo-EM data. The reference-free 2D class averages and population distributions of TRiC-NPP and TRiC-tubulin-S1 are also presented. **b-c**, Resolution estimations for TRiC-NPP and TRiC-tubulin-S1 according to the gold-standard FSC criterion of 0.143 (**b**), and local resolution evaluations (**c**).

**Extended Data Fig. 3.**
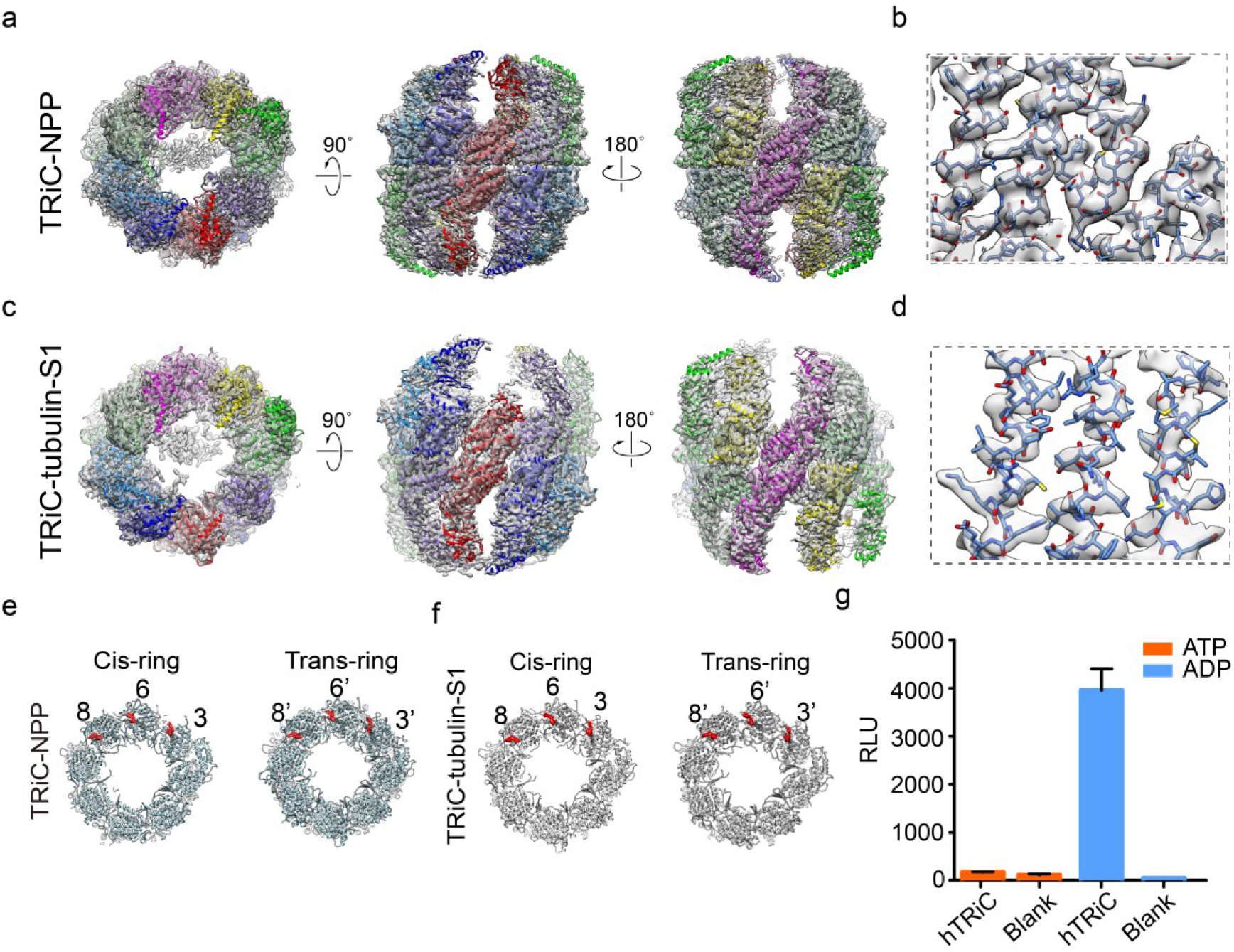
Atomic models of TRiC-NPP and TRiC-tubulin-S1. **a,c,** Atomic models of TRiC-NPP (**a**) and TRiC-tubulin-S1 (**c**) match well with the corresponding cryo-EM maps. **b,d,** Representative high-resolution structural features of TRiC-NPP (**b**) and TRiC-tubulin-S1 (**d**). **e-f**, Nucleotide occupancy statuses of TRiC-NPP (**e**) and TRiC-tubulin-S1 (**f**), with CCT3/6/8 from both rings having nucleotide density (in red) in their nucleotide pockets. **g**, ATP/ADP ratio analysis of TRiC. The relative light unit (RLU) values were measured for the sample with TRiC buffer as a blank. For all quantifications, data were plotted as mean ± SD for three independent replicates.

**Extended Data Fig. 4.**
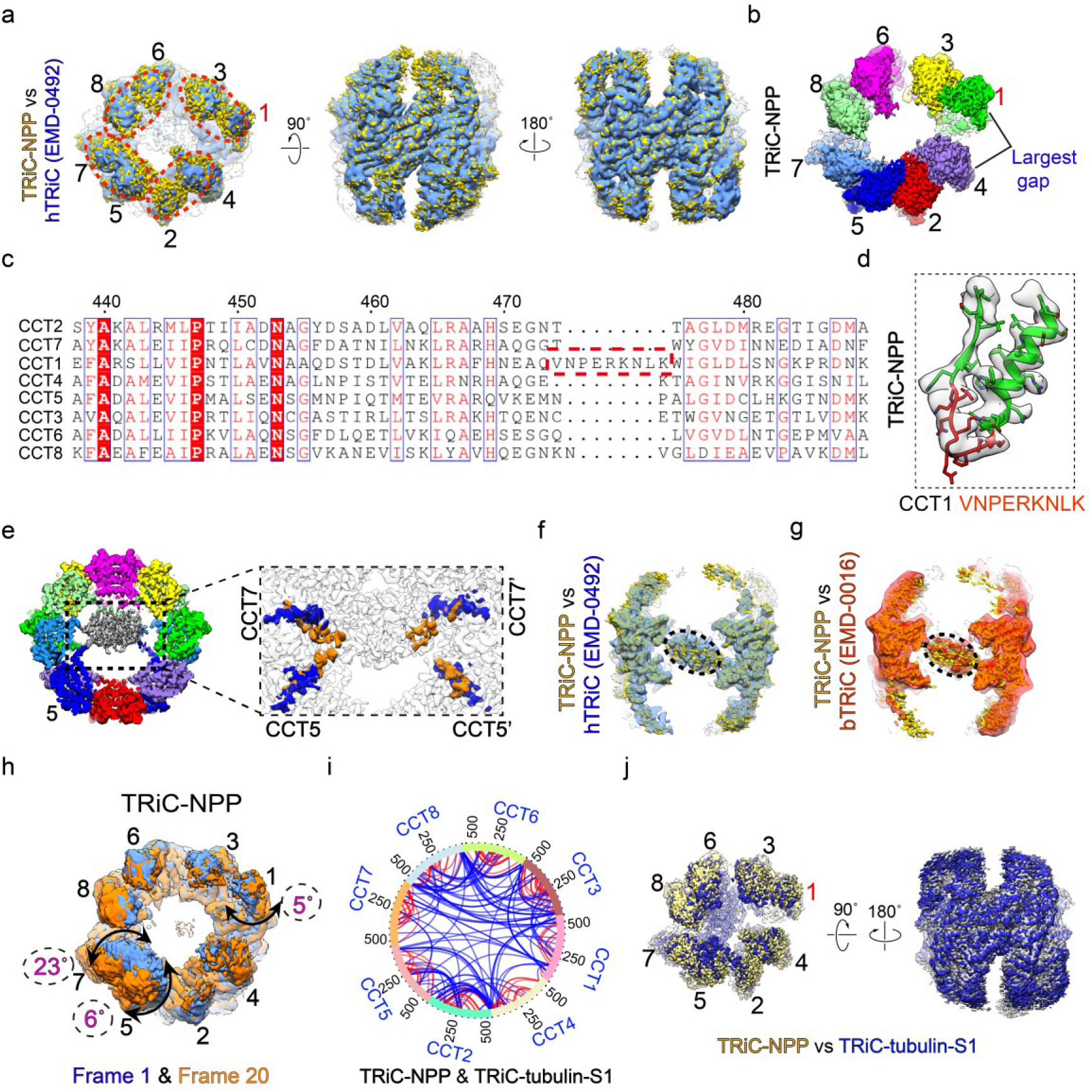
TRiC-NPP and TRiC-tubulin-S1 conformations and subunit configurations. **a**, Overlay of our TRiC-NPP map (yellow) and the reported apo hTRiC map (cornflower blue, EMDB: 0492), indicating no obvious conformational differences between the two structures. The tetramer-of-dimers pattern is indicated by dotted red circles. **b**, CCT1 and CCT4 display the largest gap between their A domains. **c-d**, Sequence alignment of all the eight subunits of hTRiC, revealing a unique insertion (indicated by red dashed frames) in the E domain of CCT1 (**c**), and the corresponding structural feature of the CCT1 E-domain insertion in TRiC-NPP (**d**). **e**, Depiction of the TRiC-NPP map showing the central tail density symmetrically contacting the N-/C-termini of CCT5/7 and CCT5’/7’ from both rings. In the magnified cut-off top view (right), only the resolved N-/C-termini of CCT5/5’ and CCT7/7’ are colored, with the C-tails in blue and N-tails in orange. **f-g**, Central slice view of the TRiC-NPP (yellow) overlaid with apo hTRiC (cornflower blue, EMDB: 0492^49^) (**f**), and with apo bovine TRiC (red, EMDB: 0016) (**g**), to show their similar tail density features (indicated by black dashed ellipsoid). **h**, Results for the 3D variability analysis (3DVA) of the TRiC-NPP dataset. This analysis suggested that CCT7/5/1 underwent a continuous outward/inward tilting motions of up to ∼23°/6°/5°, respectively. Frame 1 (cornflower blue) and frame 20 (orange), together showing the maximum extent of the conformational change, are displayed. **i**, XL-MS-derived circular plot of all XLs of the TRiC portion, with intra-subunit XLs shown in red, and inter-subunit XLs in blue. The detected cross-links within the TRiC complex fulfill the spatial geometry constrains of the linked amino acids, validating the reliability of our XL-MS data. **j**, TRiC-NPP map (yellow) overlaid with the TRiC-tubulin-S1 map (medium blue), indicating no obvious conformational differences between them.

**Extended Data Fig. 5.**
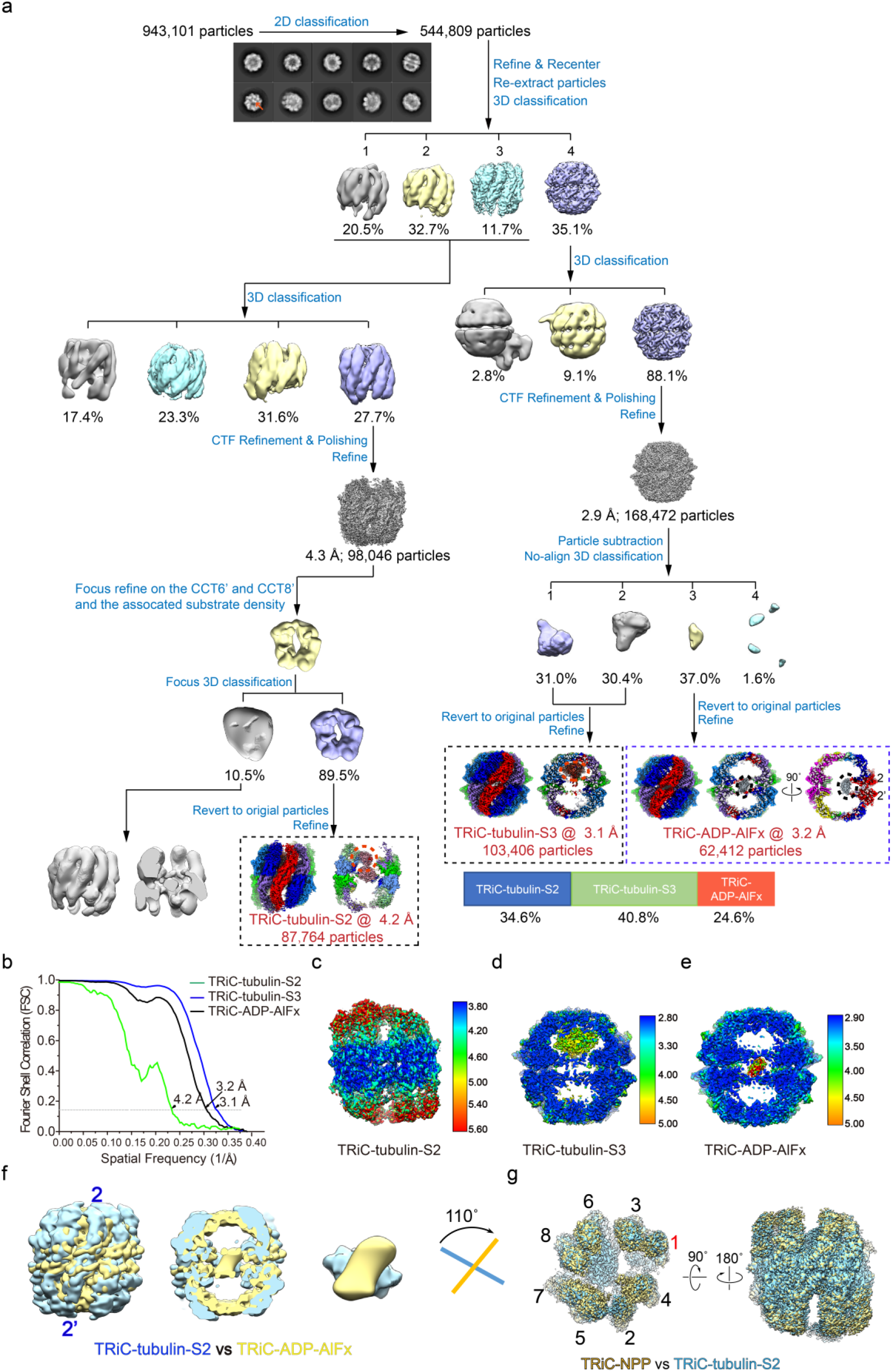
Work flow for the processing of the cryo-EM data of TRiC in the presence of ATP-AlFx. **a**, Procedure used to process the cryo-EM data of TRiC in the presence of ATP-AlFx. The reference-free 2D class averages and the population distributions of the three obtained states are also presented. Besides, central slice view of the TRiC-ADP-AlFx map revealed the tail density attached to the CCT2/2’ subunit pair. **b**, Resolution estimations of the TRiC-tubulin-S2, -S3, and TRiC-ADP-AlFx maps according to the gold-standard FSC criterion of 0.143. **c-e**, Local resolution evaluations for the S2 (**c**), S3 (**d**), and TRiC-ADP-AlFx (**e**) maps. **f**, Overlay of the TRiC-tubulin-S2 and TRiC-ADP-AlFx maps. When visualized from the typical viewing direction with the CCT2 subunit pair facing the viewer, the unstructured tail of TRiC-ADP-AlFx was observed to be rotated clockwise by about 110° relative to that of TRiC-tubulin-S2. These two maps were lowpass filtered to 8 Å for better visualization of the tail density. **g**, Overlay of the TRiC-NPP (yellow) and -S2 maps (sky blue), indicating their overall similar conformations.

**Extended Data Fig. 6.**
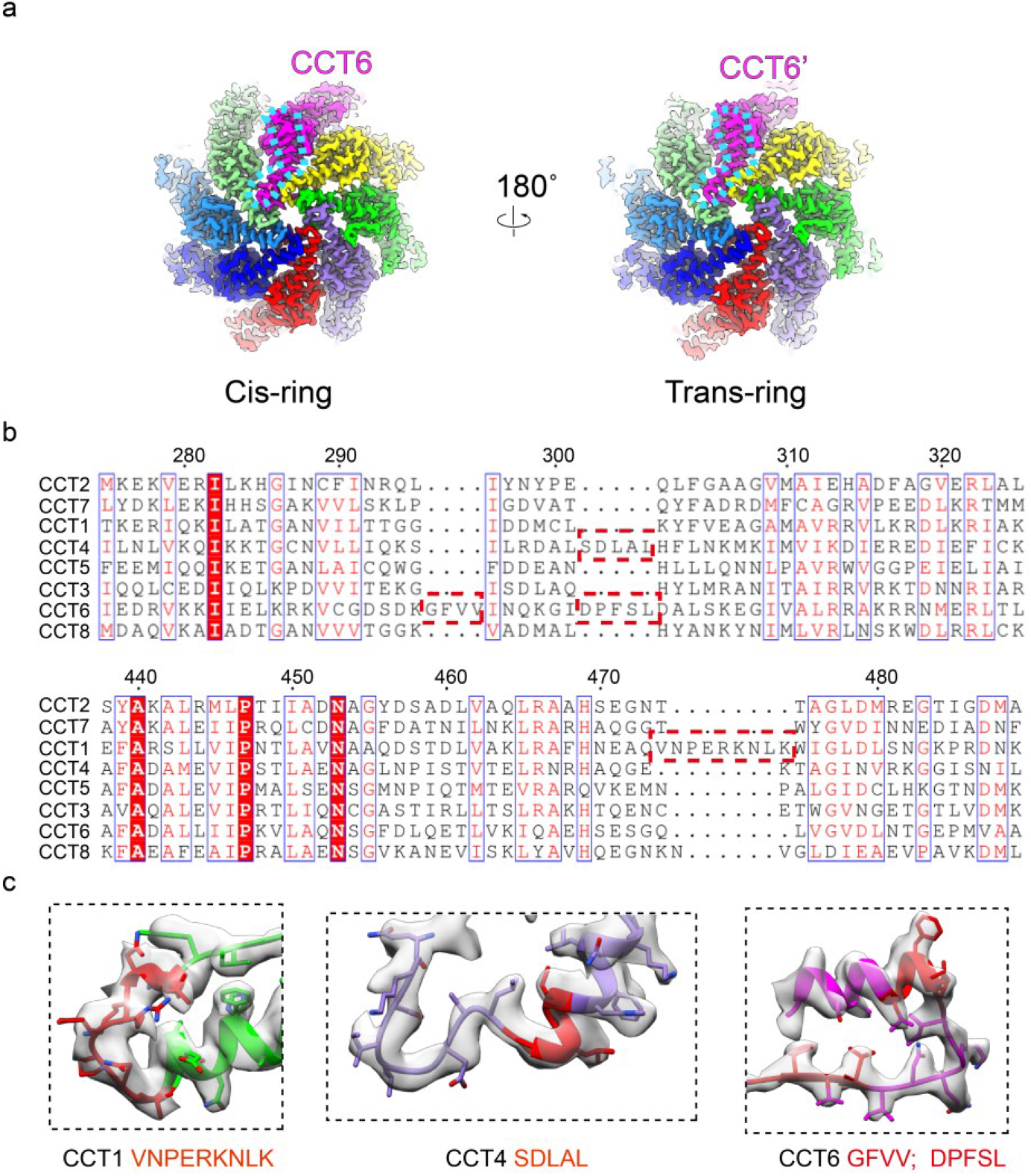
Features facilitating the subunit identification for the TRiC-tubulin-S3 map. **a**, Depictions of the S3 map in which the unique kink feature (enclosed in a dotted blue line) in the apical protrusion H8 of CCT6 was unambiguously visualized in both rings, substantiating the on-axis location of CCT6. **b**, Sequence alignment of all eight subunits of hTRiC. This alignment revealed four unique insertions (indicated by red dashed frames) in CCT1/4/6. **c**, Related structures of the four insertions (in red), all resolved in our S3 map.

**Extended Data Fig. 7.**
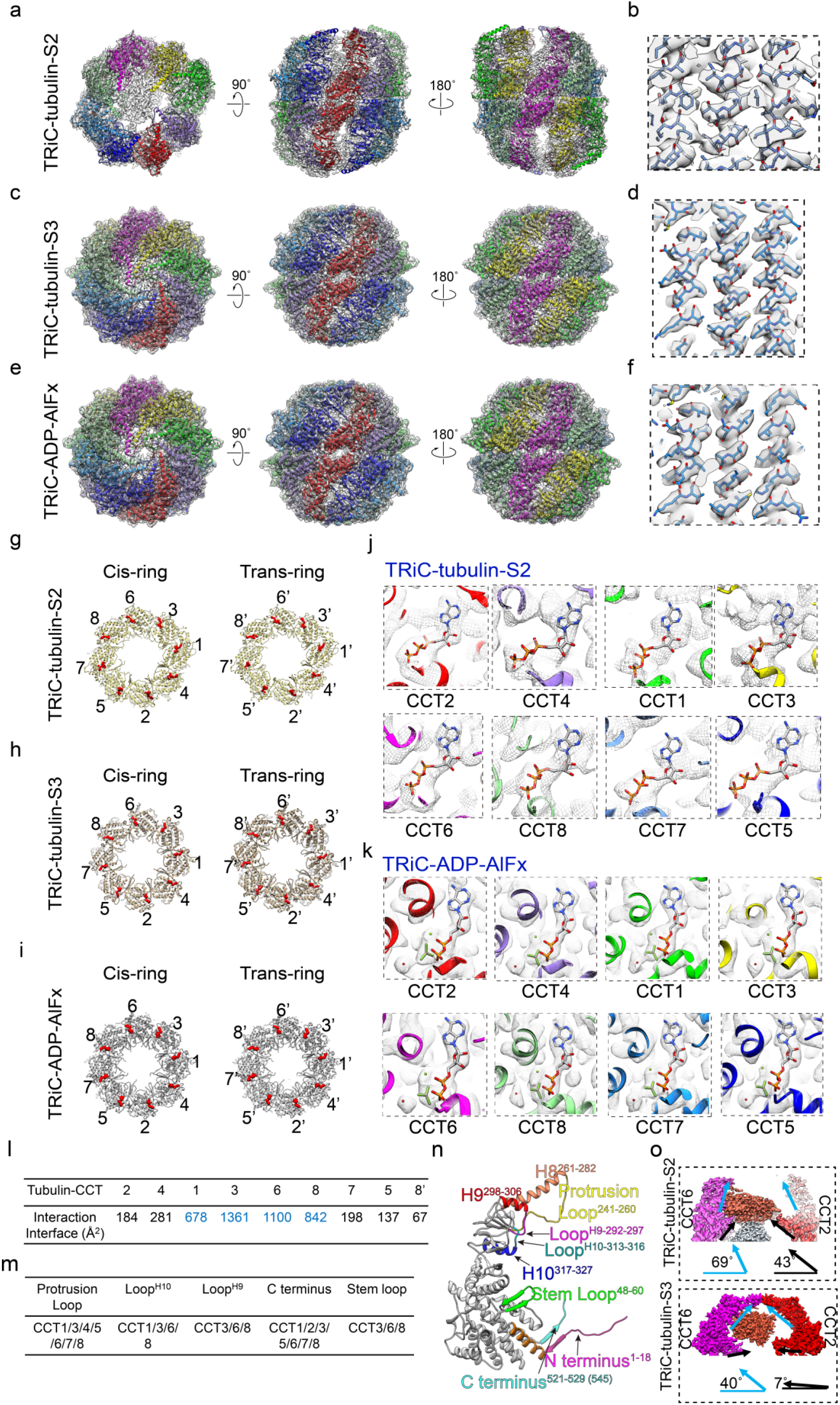
Atomic models of TRiC-tubulin-S2, -S3, TRiC-ADP-AlFx and nucleotide status in the TRiC-tubulin-S2 and TRiC-ADP-AlFx maps. **a,c,e** Atomic models of TRiC-tubulin-S2 (**a**), -S3 (**c**) and TRiC-ADP-AlFx (**e**) match well with the corresponding cryo-EM maps. **b,d,f**, Representative high-resolution structural features of TRiC-tubulin-S2 (**b**), -S3 (**d**) and TRiC-ADP-AlFx (**f**). **g-i**, Nucleotide occupancy statuses of TRiC-tubulin-S2 (**g**), -S3 (**h**) and TRiC-ADP-AlFx (**i**). **j**, Magnified view of the TRiC-tubulin-S2 nucleotide pocket region of each subunit. Every subunit appears to include a bound nucleotide density that matches the ATP model reasonably well, suggesting that the TRiC-tubulin-S2 is in the ATP-bound state. **k**, Magnified view of the TRiC-ADP-AlFx nucleotide pocket region of each subunit. Every subunit here was found to be bound with ADP-AlFx (stick model) and a magnesium ion (green ball), as well as a water molecule (red ball) in an attacking position, suggesting that each of these subunits is in the ATP-hydrolysis transition state. **l**, Interaction areas between tubulin and TRiC subunits in TRiC-tubulin-S3 as calculated using PISA. **m**, Main structural elements of TRiC involved in the interaction with tubulin in TRiC-tubulin-S3. **n**, Key structural elements of TRiC involved in the interaction with tubulin, with the residue numbers of these elements being labeled (CCT3 as a representative subunit). **o**, Depictions of a portion of the S2 and S3 maps (showing CCT6/2 subunits only), suggesting ATP-hydrolysis-induced TRiC A and E domain downward rotations. In each panel, showing is the angle between the long axis of the A (in light blue) or E domain (in black) of CCT2 and the horizontal plane for the corresponding map.

**Extended Data Fig. 8.**
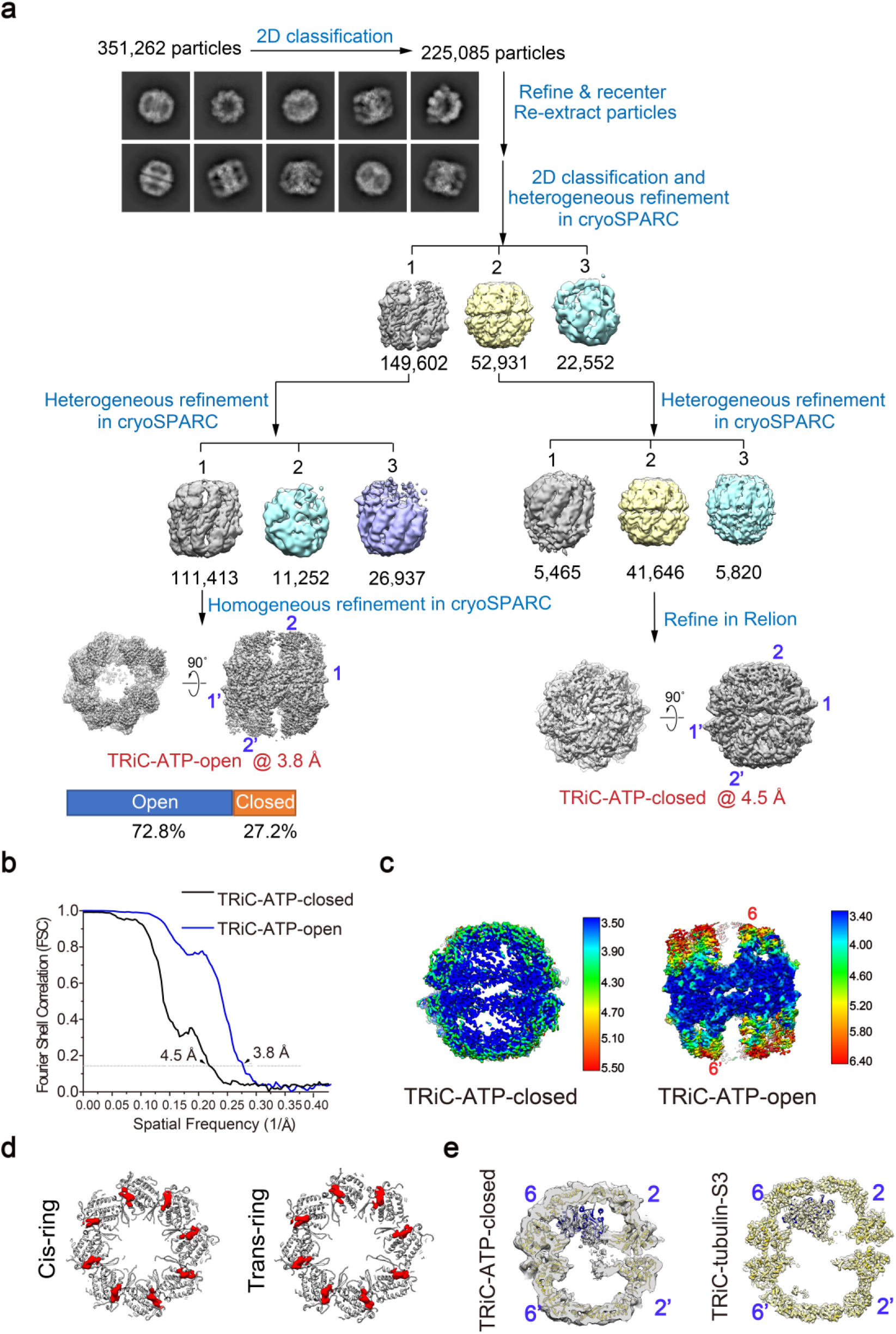
Cryo-EM analysis of TRiC in the presence of ATP. **a**, Procedure used to process the cryo-EM data of TRiC in the presence of ATP. Reference-free 2D class averages and population distributions of the closed and open conformations of TRiC (denoted as “TRiC-ATP-closed” and “TRiC-ATP-open”, respectively) are also presented. **b**, Resolution estimations for TRiC-ATP-closed and TRiC-ATP-open maps according to the gold-standard FSC criterion of 0.143. **c**, Local resolution evaluations for TRiC-ATP-closed and TRiC-ATP-open. **d**, The nucleotide occupancy status of TRiC-ATP-closed showed that all nucleotide pockets were occupied. **e**, The central slice of the TRiC-ATP-closed (left, unsharpened map) and TRiC-tubulin-S3 (right) map with fitted model of TRiC-tubulin-S3. This analysis indicates a substrate within one of the TRiC-ATP-closed chambers, in a position and orientation similar to those of tubulin observed in the TRiC-tubulin-S3 map.

**Extended Data Fig. 9.**
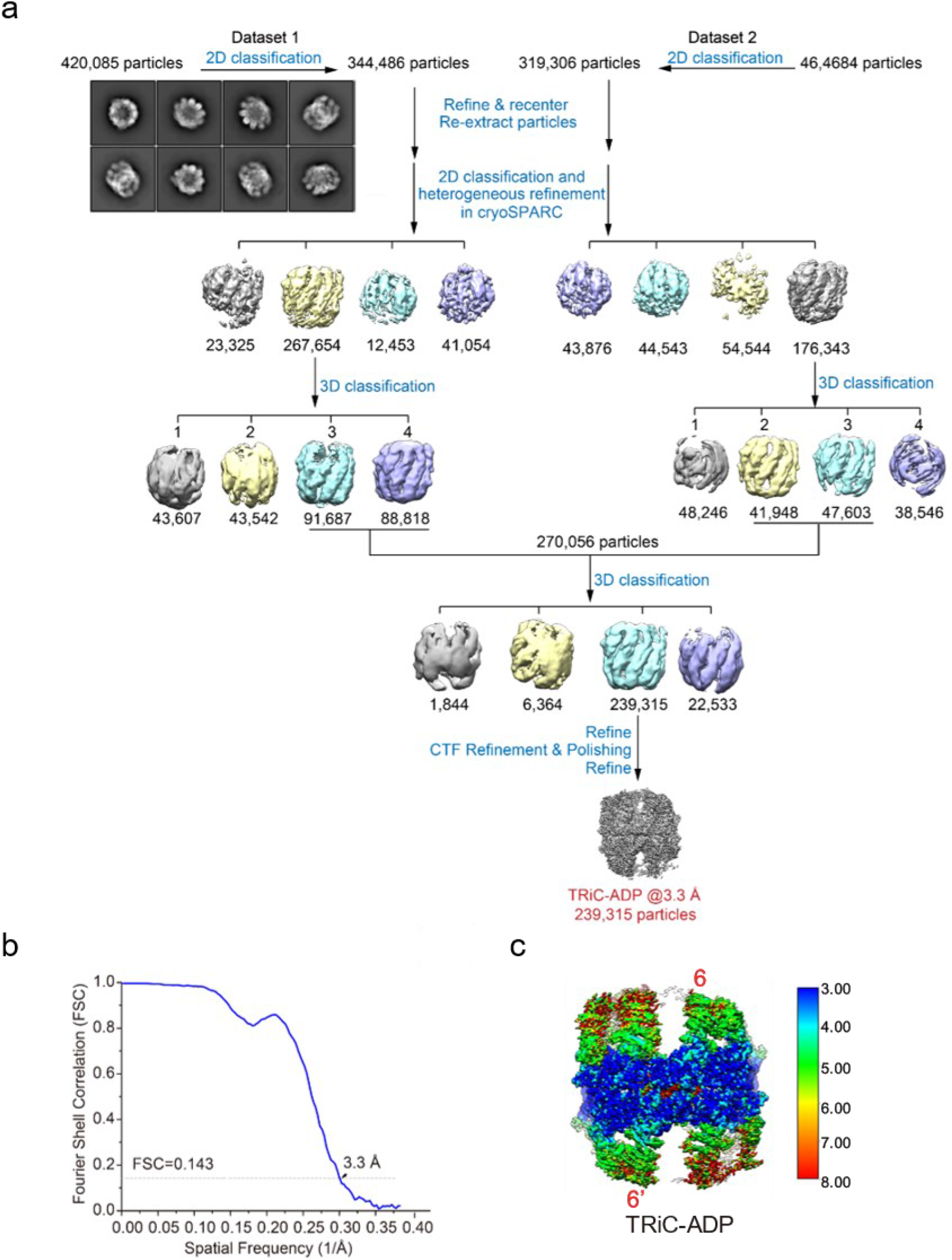
Work flow for the processing of the TRiC-ADP cryo-EM data. **a**, Procedure used to process the cryo-EM data of TRiC in the presence of ADP, with the reference-free 2D class average also shown. **b**, Estimation of the TRiC-ADP map resolution according to the gold-standard FSC criterion of 0.143. **c**, Local resolution evaluation for TRiC-ADP.

**Extended Data Fig. 10.**
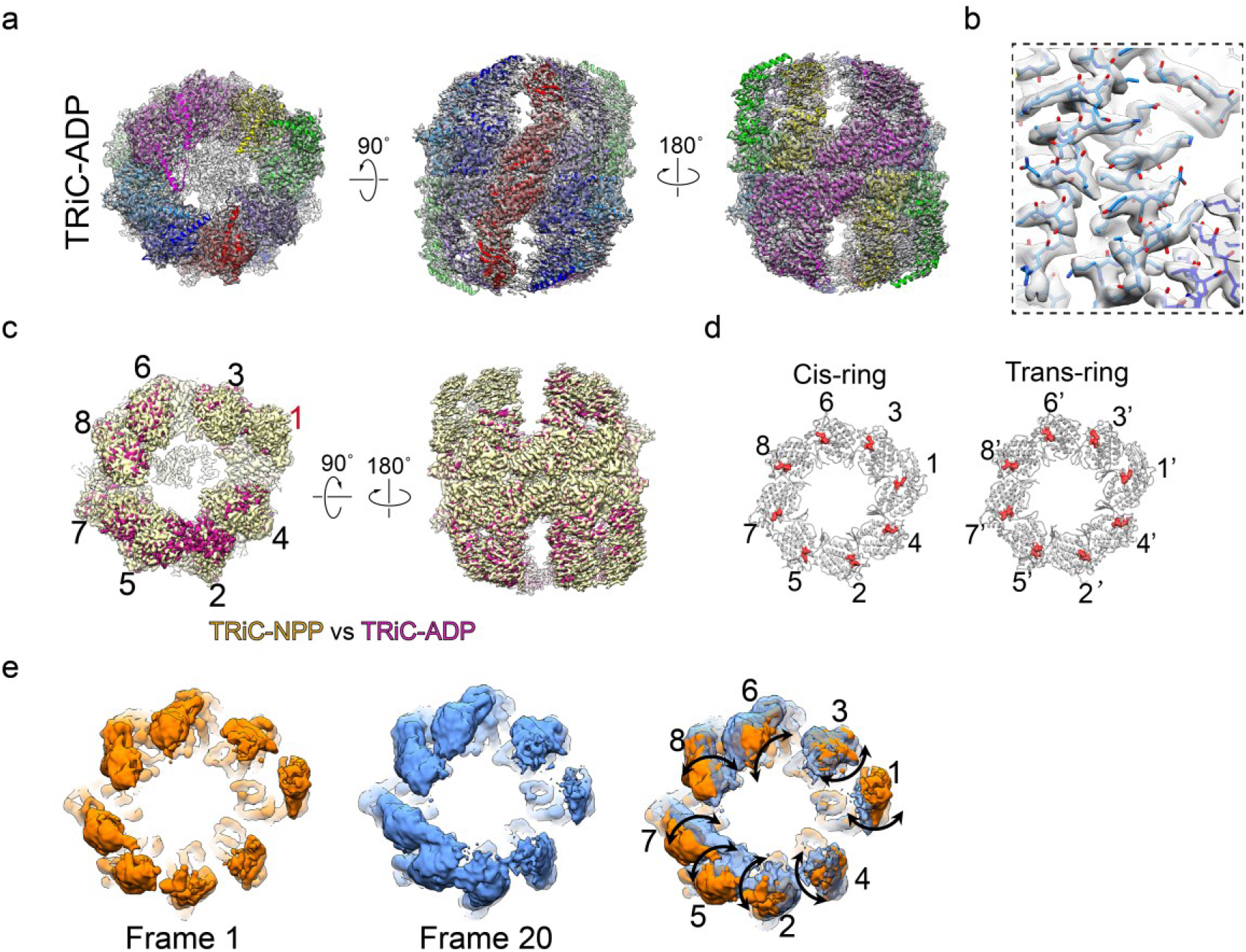
Atomic model and 3DVA result of TRiC-ADP. **a**, Atomic model of TRiC-ADP matches well with the corresponding map. **b**, Representative high-resolution structural features of TRiC-ADP. **c**, The determined nucleotide occupancy status of TRiC-ADP, showing all of the subunit nucleotide pockets were occupied. **d**, Overlay of the TRiC-NPP (yellow) and TRiC-ADP (violet red) maps, indicating no large conformational differences between them. **e**, 3DVA results of the TRiC-ADP dataset. This analysis suggested that all of the subunits underwent outward/inward tilting motions. Frames 1 (orange) and 20 (cornflower blue), together showing the maximum extent of the conformational change, are displayed here.

**Extended Data Fig. 11.**
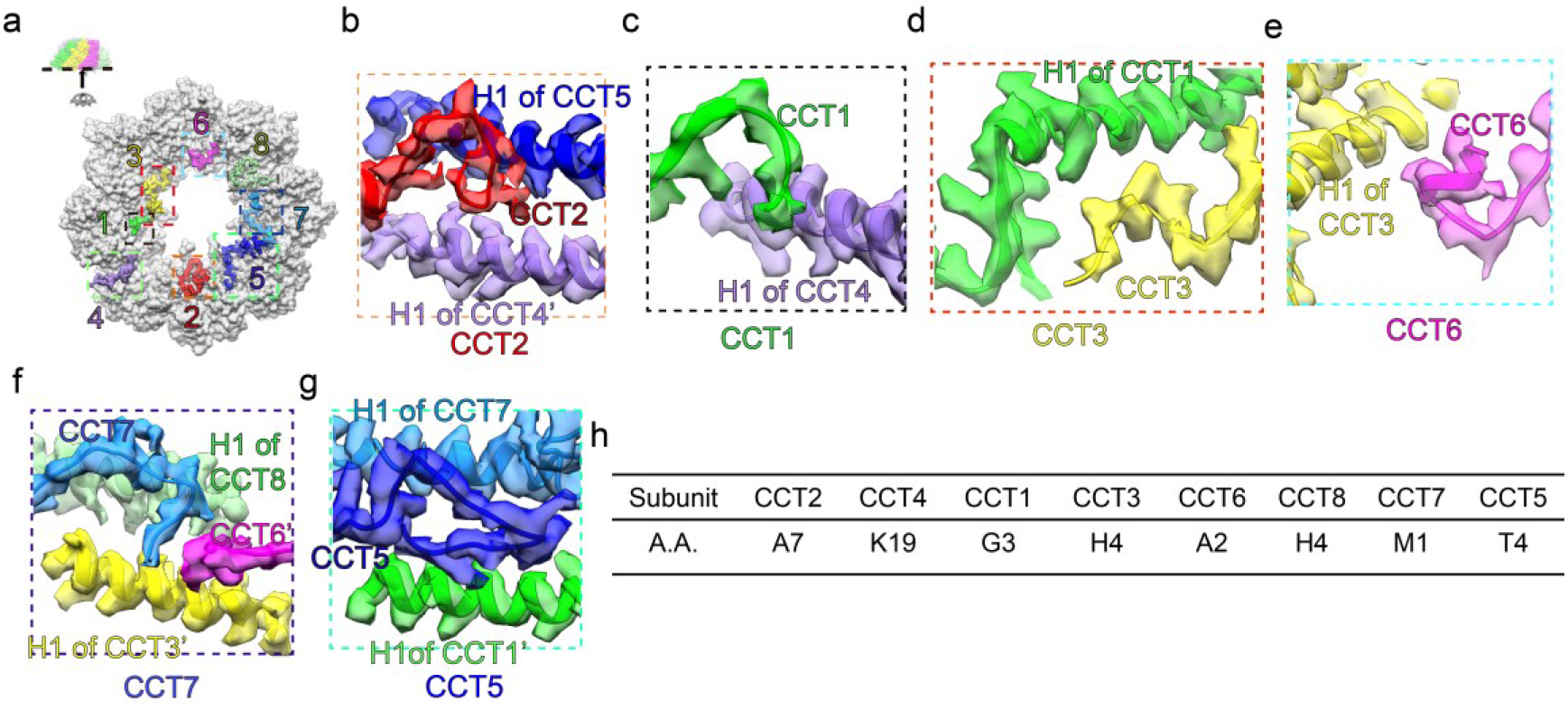
The involvement of the TRiC N-termini in the allosteric interaction network in TRiC-tubulin-S3. **a**, Locations of the resolved N-termini (in color) in the TRiC-tubulin-S3 structure. The visualization direction and visualized region are illustrated in the inset. **b-g**, Magnified views of the interaction between the N-terminus of CCT2 and helices H1 of CCT5 and CCT4’ (**b**), between the N-terminus of CCT1 and H1 of CCT4 (**c**), between the N-terminus of CCT3 and H1 of CCT1 (**d**), between the CCT6 N-terminus (there is a small 1-turn α-helix before its N-terminal β-sheet) and H1 of CCT3 (**e**), between the N-terminus of CCT7 and H1 of CCT8 and CCT3’, as well as the N-terminus of CCT6’ (**f**), and between the N-terminus of CCT5 and helices H1 of CCT7 and CCT1’(**g**). **h**, The most N-terminal amino acid residue visualized for each subunit of TRiC in the TRiC-tubulin-S3 structure.

**Supplementary Table 1.**
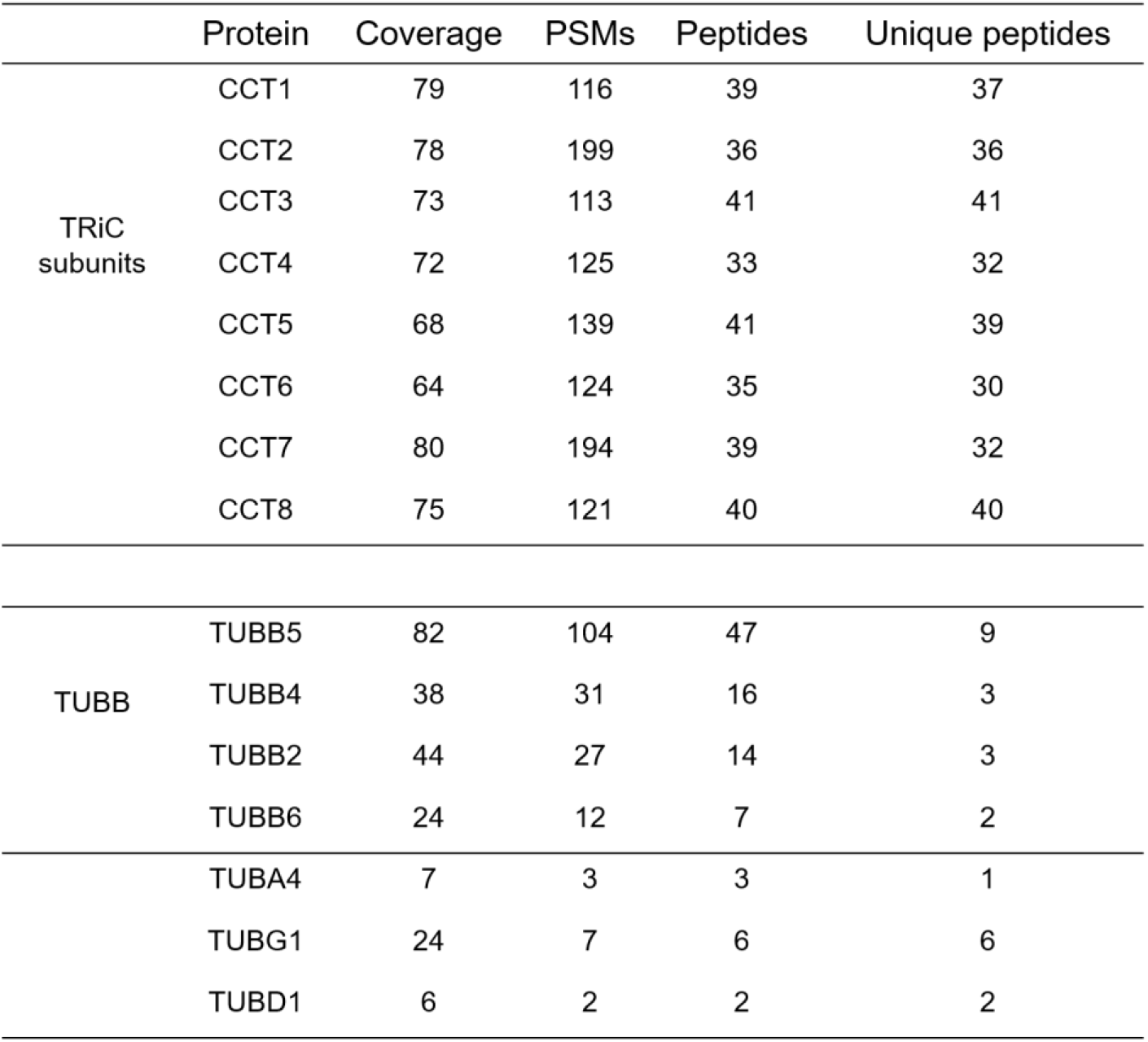
Results of mass spectroscopy (MS) analysis of endogenously purified TRiC with associated tubulin.

**Supplementary Table 2.**
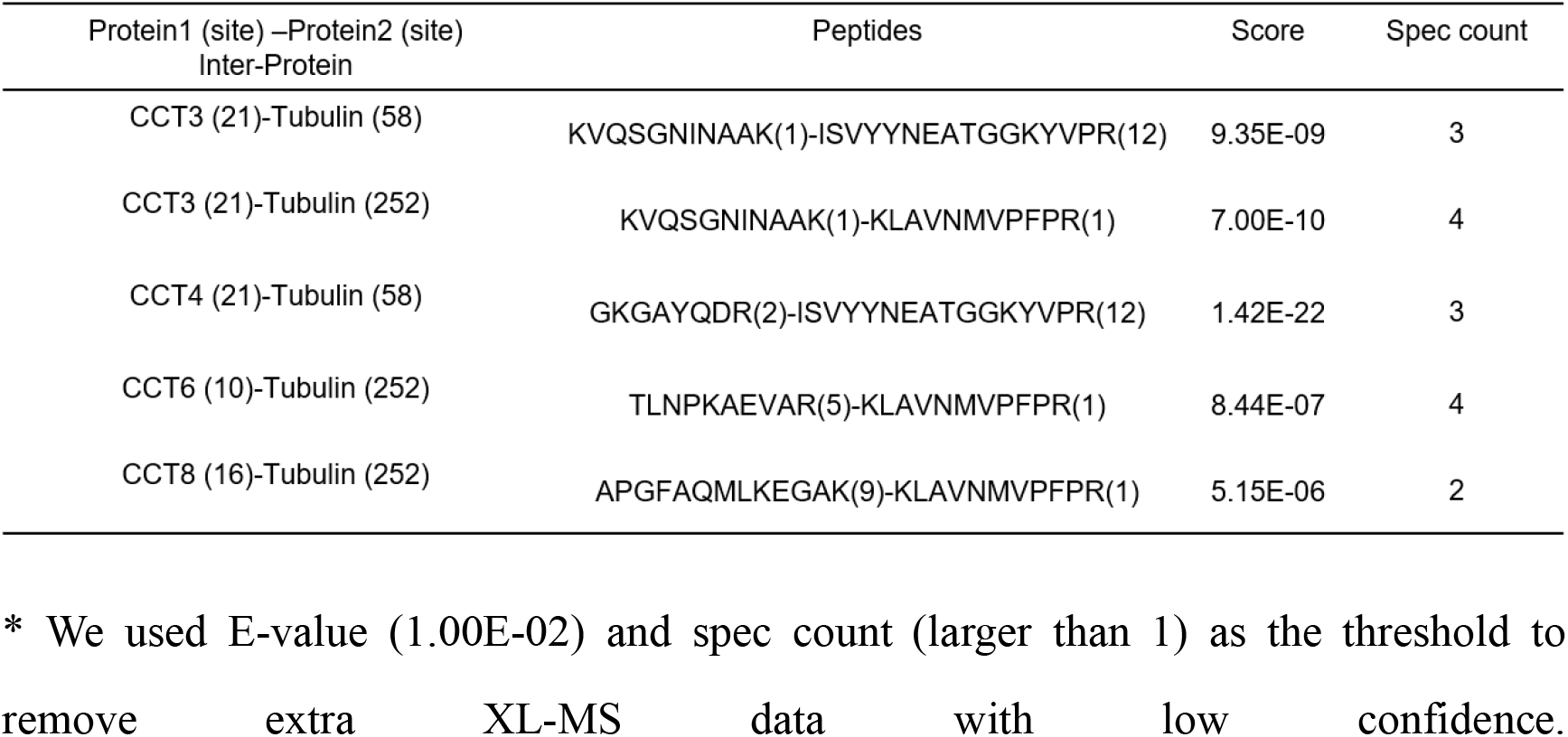
Results of XL-MS analysis of TRiC with associated tubulin.

**Supplementary Table 3.**
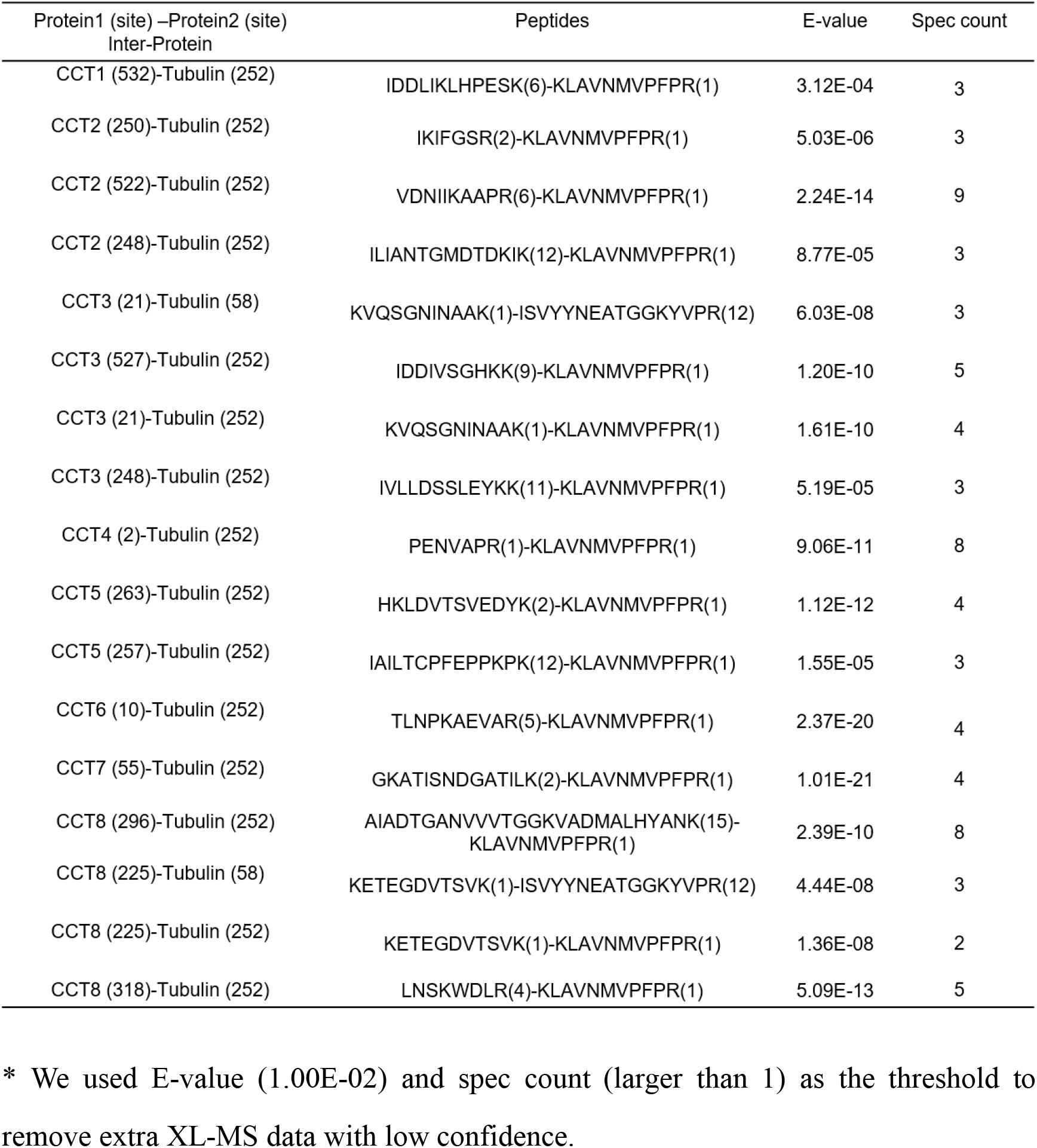
Results of XL-MS analysis of TRiC with associated tubulin in the presence of ATP-AlFx.

**Supplementary Table 4.** Interactions between tubulin and CCT3/6/8/1 detected using PISA.

|  | TRiC subunit |  | Tubulin |  | Interaction | Distance (Å) |
| --- | --- | --- | --- | --- | --- | --- |
|  | Residue | Atom | Residue | Atom |  |  |
| CCT3 | GLU 89 | [ OE1 ] | LYS 392 | [ NZ ] | Salt bridge/ H bond | 3.82 |
|  | GLU 69 | [ OE2 ] | HIS 396 | [ NE2 ] | Salt bridge | 3.32 |
|  | ARG 313 | [ NE ] | GLU 157 | [ OE2 ] | Salt bridge | 3.72 |
|  | ARG 200 | [ NH1 ] | ASP 404 | [ OD2 ] | Salt bridge/ H bond | 3.65 |
|  | ARG 316 | [ NH1 ] | GLU 407 | [ OE2 ] | Salt bridge/ H bond | 3.71 |
|  | LYS 317 | [ NZ ] | GLU 410 | [ OE1 ] | Salt bridge/ H bond | 2.52 |
|  | LYS 317 | [ NZ ] | GLU 410 | [ OE2 ] | Salt bridge | 3.87 |
|  | LYS 196 | [ NZ ] | GLU 405 | [ OE2 ] | Salt bridge | 3.90 |
|  | GLU 69 | [ OE1 ] | TRP 397 | [ NE1 ] | H bond | 3.74 |
|  | GLU 69 | [ OE2 ] | TRP 397 | [ NE1 ] | H bond | 3.64 |
|  | ARG 316 | [ NH1 ] | TYR 106 | [ OH ] | H bond | 2.48 |
|  | ARG 85 | [ NH1 ] | ARG 391 | [ O ] | H bond | 2.83 |
|  | ARG 68 | [ NH2 ] | LYS 392 | [ O ] | H bond | 3.74 |
|  | LYS 203 | [ NZ ] | GLY 400 | [ O ] | H bond | 3.30 |
|  | ARG 200 | [ NH1 ] | MET 403 | [ O ] | H bond | 3.33 |
|  | ARG 200 | [ NH2 ] | MET 403 | [ O ] | H bond | 3.49 |
|  | ASN 321 | [ ND2 ] | MET 406 | [ SD ] | H bond | 3.26 |
|  | THR 318 | [ OG1 ] | GLU 407 | [ OE2 ] | H bond | 3.03 |
| CCT6 | ASP 213 | [ OD1 ] | ARG 121 | [ NH1 ] | Salt bridge/ H bond | 3.22 |
|  | ASP 213 | [ OD1 ] | ARG 121 | [ NH2 ] | Salt bridge | 3.41 |
|  | ASP 213 | [ OD2 ] | LYS 122 | [ NZ ] | Salt bridge/ H bond | 2.45 |
|  | ARG 319 | [ NH1 ] | ASP 114 | [ OD2 ] | Salt bridge | 3.49 |
|  | ARG 319 | [ NH2 ] | ASP 114 | [ OD2 ] | Salt bridge/ H bond | 2.73 |
|  | ARG 314 | [ NH2 ] | ASP 128 | [ OD1 ] | Salt bridge/ H bond | 3.17 |
|  | ARG 318 | [ NH1 ] | GLU 157 | [ OE1 ] | Salt bridge/ H bond | 3.27 |
|  | GLU 357 | [ OE1 ] | THR 55 | [ OG1 ] | H bond | 2.84 |
|  | ARG 315 | [ O ] | ARG 121 | [ NH1 ] | H bond | 3.28 |
|  | ASP 213 | [ OD2 ] | ARG 121 | [ NH2 ] | H bond | 2.44 |
|  | TYR 239 | [ O ] | ARG 162 | [ NH1 ] | H bond | 3.48 |
|  | ALA 51 | [ N ] | ASP 74 | [ OD2 ] | H bond | 3.58 |
|  | ARG 314 | [ NH2 ] | GLU 125 | [ O ] | H bond | 2.48 |
|  | ARG 314 | [ NH1 ] | ASP 128 | [ O ] | H bond | 3.89 |
|  | ARG 217 | [ NE ] | GLU 125 | [ O ] | H bond | 3.04 |
|  | GLY 215 | [ N ] | GLU 125 | [ OE2 ] | H bond | 3.79 |
|  | GLU 238 | [ OE1 ] | LEU 130 | [ N ] | H bond | 3.75 |
|  | GLN 294 | [ NE2 ] | GLU 158 | [ OE1 ] | H bond | 2.69 |
| CCT8 | GLU 252 | [ OE1 ] | ARG 2 | [ NE ] | Salt bridge/ H bond | 3.27 |
|  | GLU 252 | [ OE1 ] | ARG 2 | [ NH2 ] | Salt bridge/ H bond | 3.26 |
|  | ARG 314 | [ NE ] | ASP 41 | [ OD2 ] | Salt bridge | 3.69 |
|  | ARG 314 | [ NH1 ] | ASP 41 | [ OD1 ] | Salt bridge | 3.51 |
|  | LYS 318 | [ NZ ] | ASP 128 | [ OD2 ] | Salt bridge/ H bond | 3.58 |
|  | GLU 252 | [ O ] | ARG 2 | [ NH2 ] | H bond | 2.98 |
|  | LYS 225 | [ NZ ] | TYR 36 | [ OH ] | H bond | 3.67 |
|  | ARG 314 | [ NH1 ] | ASP 41 | [ OD2 ] | H bond | 2.97 |
|  | ASN 316 | [ ND2 ] | ASP 41 | [ OD1 ] | H bond | 2.68 |
|  | SER 317 | [ OG ] | GLU 53 | [ OE2 ] | H bond | 2.34 |
|  | LYS 318 | [ N ] | GLU 53 | [ OE1 ] | H bond | 3.28 |
|  | TRP 319 | [ NE1 ] | GLU 53 | [ O ] | H bond | 2.66 |
| CCT1 | ARG 309 | [ NH2 ] | GLU 410 | [ OE1 ] | Salt bridge/ H bond | 3.66 |
|  | ASP 358 | [ O ] | LYS 392 | [ NZ ] | H bond | 2.94 |
|  | GLN 242 | [ NE2 ] | ASP 417 | [ OD1 ] | H bond | 3.81 |
|  | GLN 242 | [ OE1 ] | GLN 424 | [ NE2 ] | H bond | 3.55 |
|  | ARG 310 | [ NH2 ] | ASN 416 | [ O ] | H bond | 3.31 |
|  | ARG 310 | [ NH2 ] | SER 420 | [ OG ] | H bond | 2.74 |

## Supplementary Video Legends

**Supplementary Video 1 | 3D variability analysis (3DVA) of the TRiC-NPP cryo-EM data.** The 3DVA results suggested that the A/I domains of CCT1/4/2/5/7 subunits are overall relative dynamic, with those of CCT7/5/1 even displaying larger movements.

**Supplementary Video 2 | 3D variability analysis (3DVA) of the TRiC-ADP cryo-EM data.** The 3DVA results suggested that TRiC-ADP was very dynamic with all of the subunits, including the usually most stable CCT6 subunit, displaying an outward/inward tilting motion.

## Notes

### Competing Interest Statement

The authors have declared no competing interest.

### Summary of Updates

Article format updated

## Reference

1 Tam, S. et al. The chaperonin TRiC blocks a huntingtin sequence element that promotes the conformational switch to aggregation. Nature Structural & Molecular Biology 16, 1279–U1298, doi:10.1038/nsmb.1700 (2009).

2 Khabirova, E. et al. The TRiC/CCT chaperone is implicated in Alzheimer’s disease based on patient GWAS and an RNAi screen in Abeta-expressing Caenorhabditis elegans. PLoS One 9, e102985, doi:10.1371/journal.pone.0102985 (2014).

3 Pereira, J. H. et al. Mechanism of nucleotide sensing in group II chaperonins. EMBO J 31, 731–740, doi:10.1038/emboj.2011.468 (2012).

4 Hayer-Hartl, M., Bracher, A. & Hartl, F. U. The GroEL-GroES Chaperonin Machine: A Nano-Cage for Protein Folding. (2016).

5 Balchin, D., Hayer-Hartl, M. & Hartl, F. U. In vivo aspects of protein folding and quality control. Science 353, aac4354, doi:10.1126/science.aac4354 (2016).

6 Joachimiak, L. A., Walzthoeni, T., Liu, C. W., Aebersold, R. & Frydman, J. The structural basis of substrate recognition by the eukaryotic chaperonin TRiC/CCT. Cell 159, 1042–1055, doi:10.1016/j.cell.2014.10.042 (2014).

7 Balchin, D., Milicic, G., Strauss, M., Hayer-Hartl, M. & Hartl, F. U. Pathway of Actin Folding Directed by the Eukaryotic Chaperonin TRiC. Cell 174, 1507–1521 e1516, doi:10.1016/j.cell.2018.07.006 (2018).

8 Llorca, O. et al. Eukaryotic type II chaperonin CCT interacts with actin through specific subunits. Nature 402, 693–696, doi:Doi 10.1038/45294 (1999).

9 Llorca, O. et al. Eukaryotic chaperonin CCT stabilizes actin and tubulin folding intermediates in open quasi-native conformations. Embo Journal 19, 5971–5979, doi:DOI 10.1093/emboj/19.22.5971 (2000).

10 Camasses, A., Bogdanova A Fau - Shevchenko, A., Shevchenko A Fau - Zachariae, W. & Zachariae, W. The CCT chaperonin promotes activation of the anaphase-promoting complex through the generation of functional Cdc20. (2003).

11 Plimpton, R. L. et al. Structures of the Gbeta-CCT and PhLP1-Gbeta-CCT complexes reveal a mechanism for G-protein beta-subunit folding and Gbetagamma dimer assembly. Proc Natl Acad Sci U S A 112, 2413–2418, doi:10.1073/pnas.1419595112 (2015).

12 Kasembeli, M. et al. Modulation of STAT3 folding and function by TRiC/CCT chaperonin. PLoS Biol 12, e1001844, doi:10.1371/journal.pbio.1001844 (2014).

13 Trinidad, A. G. et al. Interaction of p53 with the CCT complex promotes protein folding and wild-type p53 activity. Mol Cell 50, 805–817, doi:10.1016/j.molcel.2013.05.002 (2013).

14 McClellan, A. J., Scott Md Fau - Frydman, J. & Frydman, J. Folding and quality control of the VHL tumor suppressor proceed through distinct chaperone pathways. (2005).

15 Munoz, I. G. et al. Crystal structure of the open conformation of the mammalian chaperonin CCT in complex with tubulin. Nat Struct Mol Biol 18, 14–19, doi:10.1038/nsmb.1971 (2011).

16 Jin, M. et al. An ensemble of cryo-EM structures of TRiC reveal its conformational landscape and subunit specificity. Proc Natl Acad Sci U S A 116, 19513–19522, doi:10.1073/pnas.1903976116 (2019).

17 Kalisman, N., Adams, C. M. & Levitt, M. Subunit order of eukaryotic TRiC/CCT chaperonin by cross-linking, mass spectrometry, and combinatorial homology modeling. Proc Natl Acad Sci U S A 109, 2884–2889, doi:10.1073/pnas.1119472109 (2012).

18 Wang, H., Han, W., Takagi, J. & Cong, Y. Yeast Inner-Subunit PA–NZ-1 Labeling Strategy for Accurate Subunit Identification in a Macromolecular Complex through Cryo-EM Analysis. Journal of Molecular Biology 430, 1417–1425, doi:https://doi.org/10.1016/j.jmb.2018.03.026 (2018).

19 Zang, Y. et al. Staggered ATP binding mechanism of eukaryotic chaperonin TRiC (CCT) revealed through high-resolution cryo-EM. Nat Struct Mol Biol 23, 1083–1091, doi:10.1038/nsmb.3309 (2016).

20 Zang, Y. et al. Development of a yeast internal-subunit eGFP labeling strategy and its application in subunit identification in eukaryotic group II chaperonin TRiC/CCT. Sci Rep 8, 2374, doi:10.1038/s41598-017-18962-y (2018).

21 Leitner, A. et al. The molecular architecture of the eukaryotic chaperonin TRiC/CCT. Structure 20, 814–825, doi:10.1016/j.str.2012.03.007 (2012).

22 Booth, C. R. et al. Mechanism of lid closure in the eukaryotic chaperonin TRiC/CCT. Nat Struct Mol Biol 15, 746–753, doi:10.1038/nsmb.1436 (2008).

23 Cong, Y. et al. 4.0-A resolution cryo-EM structure of the mammalian chaperonin TRiC/CCT reveals its unique subunit arrangement. Proc Natl Acad Sci U S A 107, 4967–4972, doi:10.1073/pnas.0913774107 (2010).

24 Cong, Y. et al. Symmetry-free cryo-EM structures of the chaperonin TRiC along its ATPase-driven conformational cycle. EMBO J 31, 720–730, doi:10.1038/emboj.2011.366 (2012).

25 Meyer, A. S. et al. Closing the folding chamber of the eukaryotic chaperonin requires the transition state of ATP hydrolysis. Cell 113, 369–381, doi:10.1016/s0092-8674(03)00307-6 (2003).

26 Liu, C. et al. Cryo-EM study on the homo-oligomeric ring formation of yeast TRiC/CCT subunits reveals TRiC ring assembly mechanism. BioRxiv, doi:10.1101/2021.02.24.432666 (2021).

27 Reissmann, S. et al. A gradient of ATP affinities generates an asymmetric power stroke driving the chaperonin TRIC/CCT folding cycle. Cell Rep 2, 866–877, doi:10.1016/j.celrep.2012.08.036 (2012).

28 Cuellar, J. et al. Structural and functional analysis of the role of the chaperonin CCT in mTOR complex assembly. Nat Commun 10, 2865, doi:10.1038/s41467-019-10781-1 (2019).

29 Roh, S. H., Kasembeli, M. M., Galaz-Montoya, J. G., Chiu, W. & Tweardy, D. J. Chaperonin TRiC/CCT Recognizes Fusion Oncoprotein AML1-ETO through Subunit-Specific Interactions. Biophys J 110, 2377–2385, doi:10.1016/j.bpj.2016.04.045 (2016).

30 Knowlton, J. J. et al. Structural and functional dissection of reovirus capsid folding and assembly by the prefoldin-TRiC/CCT chaperone network. Proceedings of the National Academy of Sciences 118, e2018127118, doi:10.1073/pnas.2018127118 (2021).

31 Llorca, O. et al. Analysis of the interaction between the eukaryotic chaperonin CCT and its substrates actin and tubulin. J Struct Biol 135, 205–218, doi:10.1006/jsbi.2001.4359 (2001).

32 Llorca, O. et al. The ‘sequential allosteric ring’ mechanism in the eukaryotic chaperonin-assisted folding of actin and tubulin. EMBO J 20, 4065–4075, doi:10.1093/emboj/20.15.4065 (2001).

33 Jin, M., Liu, C., Han, W. & Cong, Y. in Macromolecular Protein Complexes II: Structure and Function (eds J. Robin Harris & Jon Marles-Wright) 625–654 (Springer International Publishing, 2019).

34 Leroux, M. R. & Hartl, F. U. Protein folding: Versatility of the cytosolic chaperonin TRIC/CCT. Current Biology 10, R260–R264, doi:Doi 10.1016/S0960-9822(00)00432-2 (2000).

35 Ursic, D. & Culbertson, M. R. The Yeast Homolog to Mouse Tcp-1 Affects Microtubule-Mediated Processes. Molecular and Cellular Biology 11, 2629–2640, doi:Doi 10.1128/Mcb.11.5.2629 (1991).

36 Yaffe, M. B. et al. TCP1 complex is a molecular chaperone in tubulin biogenesis. Nature 358, 245–248, doi:10.1038/358245a0 (1992).

37 Sternlicht, H. et al. The t-complex polypeptide 1 complex is a chaperonin for tubulin and actin in vivo. Proceedings of the National Academy of Sciences of the United States of America 90, 9422–9426, doi:10.1073/pnas.90.20.9422 (1993).

38 Sullivan, K. F. Structure and Utilization of Tubulin Isotypes. Annual Review of Cell Biology 4, 687–716, doi:10.1146/annurev.cb.04.110188.003351 (1988).

39 Breuss, M. et al. Mutations in the β-tubulin gene TUBB5 cause microcephaly with structural brain abnormalities. Cell reports 2, 1554–1562, doi:10.1016/j.celrep.2012.11.017 (2012).

40 Poirier, K. et al. Mutations in the neuronal β-tubulin subunit TUBB3 result in malformation of cortical development and neuronal migration defects. Human Molecular Genetics 19, 4462–4473, doi:10.1093/hmg/ddq377 (2010).

41 Tischfield, M. A. et al. Human TUBB3 Mutations Perturb Microtubule Dynamics, Kinesin Interactions, and Axon Guidance. Cell 140, 74–87, doi:10.1016/j.cell.2009.12.011 (2010).

42 Ballatore, C., Lee, V. M. & Trojanowski, J. Q. Tau-mediated neurodegeneration in Alzheimer’s disease and related disorders. Nat Rev Neurosci 8, 663–672, doi:10.1038/nrn2194 (2007).

43 Nogales, E. & Alushin, G. M. Tubulin and Microtubule Structure: Mechanistic Insights Into Dynamic Instability and Its Biological Relevance doi:10.1016/b978-0-12-809633-8.08056-0 (2017).

44 Lin, Y. F., Tsai, W. P., Liu, H. G. & Liang, P. H. Intracellular beta-tubulin/chaperonin containing TCP1-beta complex serves as a novel chemotherapeutic target against drug-resistant tumors. Cancer Res 69, 6879–6888, doi:10.1158/0008-5472.CAN-08-4700 (2009).

45 Liu, Y. J., Kumar, V., Lin, Y. F. & Liang, P. H. Disrupting CCT-beta : beta-tubulin selectively kills CCT-beta overexpressed cancer cells through MAPKs activation. Cell Death Dis 8, e3052, doi:10.1038/cddis.2017.425 (2017).

46 Chen, J. et al. Cryo-EM of mammalian PA28alphabeta-iCP immunoproteasome reveals a distinct mechanism of proteasome activation by PA28alphabeta. Nat Commun 12, 739, doi:10.1038/s41467-021-21028-3 (2021).

47 Chaudhry, C., Horwich, A. L., Brunger, A. T. & Adams, P. D. Exploring the structural dynamics of the E.coli chaperonin GroEL using translation-libration-screw crystallographic refinement of intermediate states. J Mol Biol 342, 229–245, doi:10.1016/j.jmb.2004.07.015 (2004).

48 Clare, D. K. et al. ATP-triggered conformational changes delineate substrate-binding and -folding mechanics of the GroEL chaperonin. Cell 149, 113–123, doi:10.1016/j.cell.2012.02.047 (2012).

49 Gestaut, D. et al. The Chaperonin TRiC/CCT Associates with Prefoldin through a Conserved Electrostatic Interface Essential for Cellular Proteostasis. Cell 177, 751–765 e715, doi:10.1016/j.cell.2019.03.012 (2019).

50 Cuellar, J. et al. The Molecular Chaperone CCT Sequesters Gelsolin and Protects it from Cleavage by Caspase-3. J Mol Biol 434, 167399, doi:10.1016/j.jmb.2021.167399 (2021).

51 Punjani, A. & Fleet, D. J. 3D variability analysis: Resolving continuous flexibility and discrete heterogeneity from single particle cryo-EM. J Struct Biol 213, 107702, doi:10.1016/j.jsb.2021.107702 (2021).

52 You, L., Gillilan, R. & Huffaker, T. C. Model for the yeast cofactor A-beta-tubulin complex based on computational docking and mutagensis. Journal of Molecular Biology 341, 1343–1354, doi:10.1016/j.jmb.2004.06.081 (2004).

53 Kelly, J. J. et al. Snapshots of actin and tubulin folding inside the TRiC chaperonin. Nat Struct Mol Biol, doi:10.1038/s41594-022-00755-1 (2022).

54 Huang, H. B. et al. Physiological levels of ATP negatively regulate proteasome function. Cell Research 20, 1372–1385, doi:10.1038/cr.2010.123 (2010).

55 Guo, Q. et al. In Situ Structure of Neuronal C9orf72 Poly-GA Aggregates Reveals Proteasome Recruitment. Cell 172, 696–705 e612, doi:10.1016/j.cell.2017.12.030 (2018).

56 Machida, K., Kono-Okada, A., Hongo, K., Mizobata, T. & Kawata, Y. Hydrophilic residues 526 KNDAAD 531 in the flexible C-terminal region of the chaperonin GroEL are critical for substrate protein folding within the central cavity. J Biol Chem 283, 6886–6896, doi:10.1074/jbc.M708002200 (2008).

57 Ishino, S. et al. Effects of C-terminal Truncation of Chaperonin GroEL on the Yield of In-cage Folding of the Green Fluorescent Protein. J Biol Chem 290, 15042–15051, doi:10.1074/jbc.M114.633636 (2015).

58 Chen, D. H. et al. Visualizing GroEL/ES in the act of encapsulating a folding protein. Cell 153, 1354–1365, doi:10.1016/j.cell.2013.04.052 (2013).

59 Weaver, J. et al. GroEL actively stimulates folding of the endogenous substrate protein PepQ. Nat Commun 8, 15934, doi:10.1038/ncomms15934 (2017).

60 Hansen, W. J., Cowan, N. J. & Welch, W. J. Prefoldin-nascent chain complexes in the folding of cytoskeletal proteins. Journal of Cell Biology 145, 265–277, doi:DOI 10.1083/jcb.145.2.265 (1999).

61 Gao, Y., Vainberg, I. E., Chow, R. L. & Cowan, N. J. Two cofactors and cytoplasmic chaperonin are required for the folding of alpha- and beta-tubulin. Mol Cell Biol 13, 2478–2485, doi:10.1128/mcb.13.4.2478-2485.1993 (1993).

62 Grynberg, M., Jaroszewski, L. & Godzik, A. Domain analysis of the tubulin cofactor system: a model for tubulin folding and dimerization. BMC Bioinformatics 4, 46, doi:10.1186/1471-2105-4-46 (2003).

63 Tian, G. et al. Pathway Leading to Correctly Folded &#x3b2;-Tubulin. Cell 86, 287–296, doi:10.1016/S0092-8674(00)80100-2 (1996).

64 Hartl, F. U., Bracher, A. & Hayer-Hartl, M. Molecular chaperones in protein folding and proteostasis. Nature 475, 324–332, doi:10.1038/nature10317 (2011).

65 Rivenzon-Segal, D., Wolf, S. G., Shimon, L., Willison, K. R. & Horovitz, A. Sequential ATP-induced allosteric transitions of the cytoplasmic chaperonin containing TCP-1 revealed by EM analysis. Nat Struct Mol Biol 12, 233–237, doi:10.1038/nsmb901 (2005).

66 Ma, J. P., Sigler, P. B., Xu, Z. H. & Karplus, M. A dynamic model for the allosteric mechanism of GroEL. Journal of Molecular Biology 302, 303–313, doi:10.1006/jmbi.2000.4014 (2000).

67 Horovitz, A. & Willison, K. R. Allosteric regulation of chaperonins. Curr Opin Struc Biol 15, 646–651, doi:10.1016/j.sbi.2005.10.001 (2005).

68 Yebenes, H., Mesa, P., Munoz, I. G., Montoya, G. & Valpuesta, J. M. Chaperonins: two rings for folding. Trends Biochem Sci 36, 424–432, doi:10.1016/j.tibs.2011.05.003 (2011).

69 Dekker, C. et al. The crystal structure of yeast CCT reveals intrinsic asymmetry of eukaryotic cytosolic chaperonins. EMBO J 30, 3078–3090, doi:10.1038/emboj.2011.208 (2011).

70 Knowlton, J. J. et al. Structural and functional dissection of reovirus capsid folding and assembly by the prefoldin-TRiC/CCT chaperone network. Proc Natl Acad Sci U S A 118, doi:10.1073/pnas.2018127118 (2021).

71 Knee, K. M., Sergeeva, O. A. & King, J. A. Human TRiC complex purified from HeLa cells contains all eight CCT subunits and is active in vitro. Cell Stress Chaperones 18, 137–144, doi:10.1007/s12192-012-0357-z (2013).

72 Farr, G. W., Scharl, E. C., Schumacher, R. J., Sondek, S. & Horwich, A. L. Chaperonin-mediated folding in the eukaryotic cytosol proceeds through rounds of release of native and nonnative forms. Cell 89, 927–937, doi:10.1016/s0092-8674(00)80278-0 (1997).

73 Norby, J. G. Coupled Assay of Na+,K+-Atpase Activity. Method Enzymol 156, 116–119 (1988).

74 Polletta, L. et al. SIRT5 regulation of ammonia-induced autophagy and mitophagy. Autophagy 11, 253–270, doi:10.1080/15548627.2015.1009778 (2015).

75 Szpikowska, B. K., Swiderek Km Fau -Sherman, M. A., Sherman Ma Fau - Mas, M. T. & Mas, M. T. MgATP binding to the nucleotide-binding domains of the eukaryotic cytoplasmic chaperonin induces conformational changes in the putative substrate-binding domains. (1998).

76 Lu, S. et al. Mapping native disulfide bonds at a proteome scale. Nature Methods 12, 329–U373, doi:10.1038/Nmeth.3283 (2015).

77 Mastronarde, D. N. Automated electron microscope tomography using robust prediction of specimen movements. Journal of Structural Biology 152, 36–51, doi:10.1016/j.jsb.2005.07.007 (2005).

78 Scheres, S. H. W. Semi-automated selection of cryo-EM particles in RELION-1.3. Journal of structural biology 189, 114–122, doi:10.1016/j.jsb.2014.11.010 (2015).

79 Fernandez-Leiro, R. & Scheres, S. H. W. A pipeline approach to single-particle processing in RELION. Acta Crystallographica Section D-Structural Biology 73, 496–502, doi:10.1107/S2059798316019276 (2017).

80 Zheng, S. Q. et al. MotionCor2: anisotropic correction of beam-induced motion for improved cryo-electron microscopy. Nature methods 14, 331–332, doi:10.1038/nmeth.4193 (2017).

81 Rohou, A. & Grigorieff, N. CTFFIND4: Fast and accurate defocus estimation from electron micrographs. J Struct Biol 192, 216–221, doi:10.1016/j.jsb.2015.08.008 (2015).

82 Arnold, K., Bordoli, L., Kopp, J. & Schwede, T. The SWISS-MODEL workspace: a web-based environment for protein structure homology modelling. Bioinformatics 22, 195–201, doi:10.1093/bioinformatics/bti770 (2006).

83 Vemu, A. et al. Structure and Dynamics of Single-isoform Recombinant Neuronal Human Tubulin. Journal of Biological Chemistry 291, 12907–12915, doi:10.1074/jbc.C116.731133 (2016).

84 DiMaio, F. et al. Atomic-accuracy models from 4.5-A cryo-electron microscopy data with density-guided iterative local refinement. Nature methods 12, 361–365, doi:10.1038/nmeth.3286 (2015).

85 Adams, P. D. et al. PHENIX: a comprehensive Python-based system for macromolecular structure solution. Acta Crystallogr D 66, 213–221, doi:10.1107/S0907444909052925 (2010).

86 Emsley, P. & Cowtan, K. Coot: model-building tools for molecular graphics. Acta Crystallographica Section D-Biological Crystallography 60, 2126–2132, doi:10.1107/S0907444904019158 (2004).

87 Pettersen, E. F. et al. UCSF chimera - A visualization system for exploratory research and analysis. J Comput Chem 25, 1605–1612, doi:10.1002/jcc.20084 (2004).

88 Goddard, T. D. et al. UCSF ChimeraX: Meeting modern challenges in visualization and analysis. Protein Science 27, 14–25, doi:10.1002/pro.3235 (2018).

89 Schlee, S. et al. Prediction of quaternary structure by analysis of hot spot residues in protein-protein interfaces: the case of anthranilate phosphoribosyltransferases. Proteins 87, 815–825, doi:10.1002/prot.25744 (2019).

